# Maternal Light and Temperature Modulate Seed Longevity in Arabidopsis

**DOI:** 10.1101/2025.10.19.683148

**Authors:** S. M. Azad, D. Gil-Villar, E. Yordán, P. Mestre, J. Forment, S. Tárraga, C. González, R Niñoles, J. Gadea

## Abstract

Seed longevity is defined as the length of time a seed can preserve its germination capacity, and it is a relevant trait for disciplines from agriculture to ecology. Seed longevity is a polygenic trait influenced by genetic and environmental factors. Despite the recognized impact of climate change on many reproductive aspects of plant biology, studies focused on how these new conditions affect this trait are scarce. In this work, we describe how environmental conditions to which the plants are exposed during seed development, termed the maternal environment effect, modulate longevity of the new developed seeds. Eight *Arabidopsis thaliana* natural accessions were grown on a combination of four different environmental scenarios: 22°C or 27°C at both low-light or high-light intensity. The combined effect of both 27°C and high light conditions generated seeds with higher longevity, although the effect is accession dependent. Similarly, an anticorrelation was found between seed dormancy and seed longevity when seeds were developed at 22°C under HL conditions, highlighting the importance of the environment in determining final seed properties. Transcriptome analysis of the Bor-4 accession, whose seeds presented the highest difference in longevity between conditions, revealed a very dynamic composition of stored mRNAs modulated by the four different environmental conditions during seed development. Nearly 80% of the differentially expressed genes exhibited a combined effect when both temperature and light-intensity were altered. Results suggest that seeds subjected to higher temperature and light intensity are primed with antioxidant defences and have a higher potential to deploy raffinose family oligosaccharides, which would be critical to safeguard cellular components during the ageing process.

## Introduction

Climate change is a complex process characterized by rising levels of carbon dioxide along with increasing global temperatures, erratic precipitation events, and higher light exposure due to cloud cover reduction. The pace of these changes has accelerated significantly in the last decade, and climate models forecast that global temperatures will continue to rise significantly over the next 50 years, with increases of 2-4°C by 2100 (Lee et al. 2023). Increases in ambient temperature have reduced global yields of major staple crops (Zhao et al. 2017). This underscores the necessity to investigate the impact of environmental changes on all aspects of plant biology to ensure future food security and sustainable agricultural practices.

Plants have evolved multiple mechanisms to perceive environmental cues. Body form, cellular physiology and gene expression are adjusted to new conditions, potentially improving the chances of seedling establishment, growth and plant survival. Mild elevation of temperature of about 5°C triggers a plethora of adaptations (which are different from the ones activated under severe heat stress) (Kan et al. 2023). These adaptations, termed thermomorphogenesis, help plants to acclimate to a warmer suboptimal environment, and to mitigate their negative effects. In Arabidopsis, readjustments lead to hypocotyl elongation, thermonasty, petiole elongation and reduced stomatal density (Koini et al. 2009, Crawford et al. 2012, Mizutani and Kanaoka 2018, Casal and Balasubramanian 2019). Similarly, plants acclimated to high light intensity modulate leaf architecture, chloroplast structure, composition of the photosynthetic electron transport chain, and other mechanisms regulating light efficiency (Weston et al. 2000, Ballottari et al. 2007, Albanese et al. 2016). High irradiation and temperature often occur simultaneously in nature, and the intertwining of both signals generates a complex scenario that differs from the impact of individual ones, at least under stress conditions (Mittler 2006, Zandalinas et al. 2020, Pascual et al. 2022, Zandalinas and Mittler 2022). In Arabidopsis, the combination of these stresses at moderate intensities associates with acclimation responses (Balfagón et al. 2019, Balfagón et al. 2022). For example, an increase in jasmonic acid (JA) fosters plant survival and is associated with elevated expression of the transcription factors (TFs) ZAT6 and ZAT10, and antioxidative enzymes (Balfagón et al. 2019).

Seed performance - the ability of seeds to successfully germinate, grow vigorously, and produce healthy plants-is critical for the plant life cycle. Environmental signals, including temperature, photoperiod, light intensity levels, and nutritional status, are known to shape seed traits (Penfield and MacGregor 2017, Zinsmeister et al. 2020, Klupczyńska and Pawłowski 2021). Multiple biochemical and molecular adaptations are orchestrated during seed maturation to generate seeds more adapted to new environments, although morphological changes are minor due to constraints in the seed structure. Embryogenesis is not affected when plants are grown up to 27°C (Malabarba et al. 2021), but seeds are smaller if parentals are grown in summer conditions (higher irradiance and temperature) (Andalo et al. 1999). Seeds developed at warmer temperatures are generally less dormant and have higher germination capacity than those that were developed at lower temperatures, correlating with lower abscisic acid (ABA) and higher gibberellin (GA) levels in those seeds (Chiang et al. 2011, Kendall et al. 2011, Chen et al. 2014, Springthorpe and Penfield 2015). Light intensity during seed development also influences dormancy acquisition by affecting the ABA/GA hormone ratio and gene expression, with lower light often leading to increased seed dormancy (Liu et al. 2025).

Seed longevity, the ability of a seed to remain viable for extended periods, is a critical adaptive trait. The oxidative damage generated in seed tissues during ageing leads to a progressive deterioration of seed quality, culminating in the inability of the embryo to germinate. Mechanisms to resist deterioration are acquired during the final stages of seed development (Righetti et al. 2015), characterised by the accumulation of protective compounds, such as antioxidants, late embryogenesis abundant (LEA) proteins, heat-shock proteins (HSPs) and raffinose family oligosaccharides (RFOs), as well as repair mechanisms that will be activated during seed imbibition to fix the damage that occurred to cellular components during storage (Rajjou and Debeaujon 2008). Increases in light intensity and temperature can be detrimental to seed longevity, as metabolism is intensified and produces more reactive oxygen species, but acclimation and protective mechanisms can be enhanced and seed longevity modulated so that damage is repaired, and seedling emergence can overcome periods of unfavourable conditions. The underlying mechanisms governing this process are unknown (Zinsmeister et al. 2020).

A wide range of variability in seed longevity exists among *Arabidopsis thaliana* natural accessions when grown under standard conditions (Renard et al. 2020), but only a few studies have investigated the effects of mild temperature or light intensity increases on this trait. A study in the widely used reference Columbia (Col-0) and Landsberg (Ler) accessions uncovered a prominent contribution of the genetic background (He et al. 2014) and revealed these cues as two of the most discriminative parental environments determining longevity. In that study, increased seed longevity was observed when seeds developed at warm temperature (25°C) or high light conditions (300 μmol m^-2^ s^-1^). This result contrasts with that reported by other study performed in Col-0, where seeds developed at 25°C presented a decrease in seed longevity (Malabarba et al. 2021), an effect which was exacerbated at 27°C. The behaviour of other natural accessions under climate change environments has not been explored.

The link between seed dormancy and longevity also remains elusive, as both positive and negative correlations between these two traits have been reported. Seeds from mutant lines in seed coat components (Debeaujon et al. 2000, De Giorgi et al. 2015) or ABA biosynthesis or signalling (Clerkx et al. 2003, Nakashima et al. 2009) exhibit a positive correlation, whereas studies using natural allelic variants of the *DOG1* dormancy gene (Nguyen et al. 2012) mutants of the auxin biosynthesis pathway (Pellizzaro et al. 2020) or the Col-0 and Ler natural accessions (He et al. 2014) revealed an inverse relationship between both traits. Most of these studies were performed on seeds developed in standard conditions. The relationship between both traits under climate change conditions is essential to get a better understanding of this trade-off.

In this study, the effect of parental light intensity and temperature on seed dormancy and seed longevity is evaluated in eight natural accessions of Arabidopsis. Our results reveal a complex interplay between genotype and environment, underscoring the complexity of identifying general trends or single variables as solely responsible for observed phenotypes in these agronomic traits. Characterisation of the stored mRNA composition for the Bor-4 accession, which presents a high environmental plasticity for seed longevity, uncovers high levels of transcripts from antioxidant genes accumulated in seeds developed at high light intensity and 27°C, highlighting the importance of protection against oxidative stress as a primary defence mechanism in mitigating seed aging-related deterioration.

## Materials and Methods

### Plant material and Growth conditions

*Arabidopsis thaliana* accessions were obtained from the Nottingham Arabidopsis Stock Centre (NASC).All accessions were grown at 22°C or 27°C in growth chambers with a 16-hour light and 8-hour dark photoperiod and 70–75% relative humidity (RH), in two light source conditions: 90 μmol photons/m²·s as low light (LL) and 300 μmol photons/m²·s as high light (HL).

### Seed assays

Controlled-deterioration treatments were conducted by storing seeds at 37°C and 75% RH, maintained with a NaCl-saturated solution in a desiccator, for 18, 25, 32, and 39 days. For germination tests, seeds were sterilized by soaking in 70% ethanol containing 0.1% Triton X-100 for 15 min, followed by three rinses with sterile water. Stratification of seeds for three days at 4°C in the dark was performed before sowing them on Murashige and Skoog medium containing 1% (wt/vol) sucrose, 10 mM 2-(N-morpholino) ethanesulfonic acid (MES buffer), and 0.9% agar with an adjusted pH of 5.7. Germination (radicle emergence) was scored after seven days. Germination rates were evaluated to determine the time for 50% germination (P50). A quadratic equation, y = ax² + bx + c, was fitted to the germination data for each treatment. The P50 value was found by solving for x, with y set to 50. a, b, and c coefficients were obtained from the curve. For seed dormancy, some seeds were tested for germination immediately after harvest (day 0). The remaining ones were kept in a desiccator at 22°C, where relative humidity was controlled by a saturated solution of Calcium Nitrate (Ca (NO3)2). Samples were taken on days 4, 7, 14, 21, 28, and 42 (until 100% of the seeds germinated in all accessions). For this assay, germination was tested by sowing seeds on plates containing two layers of autoclaved Whatman filter paper moistened with 4 ml of sterile. For all assays, the mean of three biological replicates, each containing 50 seeds, was used to determine the percentage of germination. To calculate DSDS50 (days of seed dry storage required to reach 50% germination), the germination percentage was plotted against the days of storage, and a curve was fitted using a logistic regression model.

### RNA extraction and qPCR

Total RNA was extracted from dry seeds as in Oñate-Sánchez and Vicente-Carbajosa 2008). The Maxima first-strand cDNA synthesis kit (Thermo Fisher Scientific) was used to reverse-transcribe 1 μg of RNA, according to the manufacturer’s instructions. The PyroTaq EvaGreen qPCR Mix Plus (ROX; Cultek S.L.U.) was used in a 20 μl total volume for quantitative real-time polymerase chain reactions (qRT-PCR) utilizing an Applied Biosystems 7500 Real-Time PCR System (Thermo Fisher Scientific). The mean of three replicates is used. A heat-dissociation curve (from 60 to 95°C) was used to test the specificity of PCR amplification. The ASAR1 gene expression (*AT4G02080*) was chosen as internal standard (Czechowski et al. 2005). The comparative ΔCt technique was used to determine relative mRNA abundance. Supplemental Table 9 contains a list of primers for qRT-PCR experiments.

### RNAseq analysis

Twenty-million 150nt paired-end reads per library were sequenced. After adaptor removal and low-quality trimming of raw reads with cutadapt (Martin, 2011), clean reads were quality assessed with FastQC (Andrews, 2010) and mapped to the TAIR10 Arabidopsis thaliana genome assembly using STAR (Dobin et al, 2013). The number of uniquely mapped read counts was also obtained using STAR with the Araport11 annotation. Differential expression analysis was done with DESeq2 (Love et al, 2014). Significative genes (p-adjusted <0,05) were used for functional analysis using AgriGOv2 (Tian et al, 2017). Redundant GO terms were removed and remaining GO terms were visualised using ReviGO (Supek et al, 2011).

### Metabolite analysis

Primary metabolites extraction was performed by homogenising samples in liquid nitrogen; then 100% methanol with 60 μl of internal standard (ribitol at 0.2 mg/ml in H2O) was added. The mixture was incubated for 15 minutes at 70°C and then centrifuged for 10 minutes at 14,000 rpm. The supernatant was transferred to a new tube, to which 750 μl of chloroform and 1500 μl of H2O were added. Samples were centrifuged for 15 minutes at 14,000 rpm and 150 μl of supernatant was dried in a speed-vac for 3 hours. To derivatise, the dry residues were resuspended in 40 μl of 20 mg/ml methoxyamine hydrochloride in pyridine and incubated for 2 hours at 37°C. Next, 70 μl of MSTFA-Mix (1 ml of N-Methyl-N-(trimethylsilyl) trifluoroacetamide + 2 μl of FAM-Mix) was added, incubated for 30 minutes at 37°C and transferred to a measuring vial. For gas chromatography and mass spectrometry analysis, 2 μl of each sample was injected in split mode (1:10) into a 6890N gas chromatograph (Agilent Technologies Inc., Santa Clara, CA) coupled to a Pegasus 4D TOF mass spectrometer (Leco, St. Joseph, MI). Gas chromatography was performed using a BPX35 column (30 m x 0.32 mm x 0.25 μm) (SGE Analytical Science Pty Ltd., Australia) with helium as the carrier gas at a constant flow of 2 ml/minute. The liner was set to a temperature of 230°C. The oven programme was set to 85°C for 2 minutes and a ramp of 8°C/minute up to 230°C. The chromatograms and mass spectra were analysed using Chromatof software (LECO, St. Joseph, MI) and the compounds of interest were identified by comparison with previously spiked pattern spectra.

### Detection of mitochondrial superoxide

To detect mitochondrial reactive oxygen species (ROS), MitoSOX™ Red Mitochondrial Superoxide Indicators (M36007, Invitrogen) was used. Both dry and pre-hydrated seeds (15 hours in water prior to dissection) were used. Embryos were stained with 5 μM MitoSOX™ Red (in 1X PBS) for 30 minutes at 37°C in the dark, followed by three washes with 1X PBS. Samples were mounted on microscope slides, and images were acquired and visualized using a Zeiss LSM780 laser-scanning confocal microscope equipped with a White Light Laser (WLL) (Carl Zeiss), using appropriate excitation and emission settings (482 nm excitation; 552–631 nm detection). All images were acquired under similar microscope acquisition settings. Z-stacks were obtained and processed into maximum intensity projection images using ImageJ software.

## Results

### Parental environment modulates seed longevity in an accession-specific manner

To investigate the effects of parental light intensity and temperature on seed longevity, we applied a combination of these cues to eight natural accessions of *Arabidopsis thaliana*. Accessions were selected based on their seed longevity behaviour when plants were grown at 22°C under artificial light (150μmol m^-2^ s^-1^) (Niñoles et al. 2022): 3 “long-lived” accessions with higher total germination (TG) (Da-0, Mt-0 and Chat-1), and 4 “short-lived” ones with lower TG than Col-0 after 18 months of ambient ageing (Tou-A1-67, Par-4, Bor-4 and Ler). Col-0 was also used for comparison. Plants were grown at low light (LL, 90μmol m^-2^ s^-1^), high light (HL, 360μmol m^-2^ s^-1^), low temperature (22°C) and high temperature (27°C), in four combinations (22°C/LL; 22°C/HL; 27°C/LL; 27°C/HL). Flowering was accelerated in plants grown at 27°C (Casal and Balasubramanian 2019), but no morphological signs of severe heat or light stress (Muhlenbock et al. 2008, Kan et al. 2023) were observed in the plants. Seed yield was generally lower when plants were grown at 27°C, especially for some accessions. Seeds were harvested and seed longevity assayed using controlled deterioration treatment (CDT). All batches exhibited more than 90% germination before ageing (Supplemental Table 1). Seeds from our “basal” conditions (22°C/LL) followed the same trend as in Niñoles et al. 2022, with “long-lived” accessions producing seeds with higher longevity than those catalogued as “short-lived” (Table 1).

**Table 1:** P50 values for *Arabidopsis thaliana* accessions under four environmental conditions. Values are expressed as Mean ± SD. LL: low-light conditions. HL: high-light conditions. SD: standard deviation.

|  | 22°C/LL | 22°C/HL | 27°C/LL | 27°C/HL |
| --- | --- | --- | --- | --- |
| **Da-0** | 20.9 ± 2.2 | 35.7 ± 1.3 | 23.8 ± 2.3 | 31.4 ± 0.4 |
| **Mt-0** | 20.2 ± 2.6 | 28.4 ± 0.4 | 33.1 ± 0.4 | 28.9 ± 3.7 |
| **Chat-0** | 24.4 ± 1.2 | 28.8 ± 1.0 | 30.2 ± 0.9 | 26.9 ± 1.7 |
| **Col-0** | 18.8 ± 1.9 | 29.2 ± 1.2 | 25.9 ± 3.1 | 31.3 ± 1.1 |
| **Ler** | 10.3 ± 1.2 | 24.5 ± 1.8 | 17.0 ± 1.7 | 29.2 ± 0.7 |
| **Tou-A1-67** | 10.2 ± 0.3 | 09.9 ± 0.9 | 08.6 ± 0.5 | 13.2 ± 0.4 |
| **Par-4** | 11.2 ± 1.6 | 12.9 ± 0.7 | 10.4 ± 1.0 | 28.5 ± 1.1 |
| **Bor-4** | 09.3 ± 0.0 | 23.2 ± 1.8 | 13.9 ± 3.0 | 34.5 ± 1.4 |

Overall, mild increases of light intensity or temperature had a positive impact on seed longevity in all accessions, although global increments (defined as from 22°C/LL to 27°C/HL) and the relative contribution of both cues were highly dependent on the genetic background. For example, the global increment ranged from 11% in Chat-0 to 316% in Bor-0. “Short-lived” accessions displayed significantly higher percentage of increase than “long-lived” ones, with increases of 154.9% in Par-4, 178.8% in Ler, and 316% in Bor-4. Light intensity had a more pronounced contribution than temperature. For six accessions (Da-0, Col-0, Bor-4, Ler, Tou-A1-67 and Par-4), maximal increments for individual cues were observed at HL while maintaining temperature constant (at 22°C for Da-0, Col-0, Bor-4 and Ler, and at 27°C for Tou-A1-67 and Par-4). Combined cues (as comparing 22°C/LL against 27°C/HL) resulted in higher longevity only for three of the accessions (Col-0, Ler and Bor-4). Global increments were neither additive nor synergic (except for Bor-4), suggesting complex genotype-specific cross-talks between the two signals. For example, for the Par-4 accession, HL causes a high increase in longevity only if the plant temperature is 27°C. For Ler, in contrast, the light effect is more relevant when plants are growing at 22°C, whereas Bor-4 seeds exhibit a temperature-independent light influence. Similarly, for Par-4, a temperature-effect on seed longevity appears only in HL seeds, whereas in Mt-0, the effect is more relevant at LL conditions (Figure 1). In summary, mild increases in light intensity or temperature generate seeds with higher longevity, with effects more relevant in accessions less prepared to combat deterioration.

**Figure 1:**
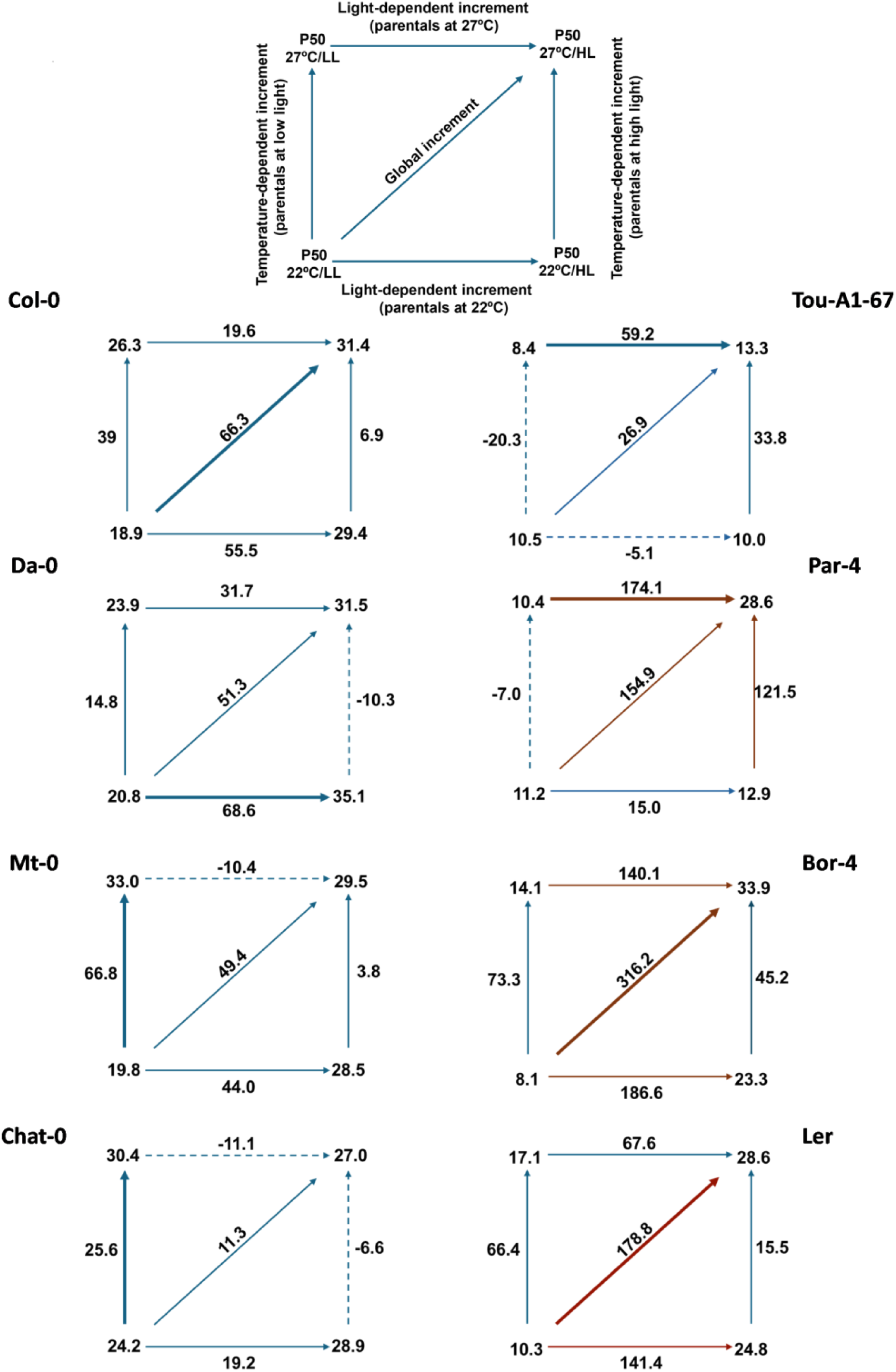
Average P50 increments (in percentage) observed for seeds of *Arabidopsis thaliana* accessions grown under different environmental conditions. Solid lines: positive increments; dashes lines: negative increments; thick line: biggest increment per accession. LL: low-light conditions. HL: high-light conditions

### Parental environment modulates seed dormancy in an accession-specific manner

Dormancy was assayed in seeds harvested from the previous experiment. As reported in other studies (Huang et al. 2018), seeds from plants grown at higher temperature exhibit lower dormancy. DSDS50 comparisons could only be calculated from five out of the eight accessions when they were grown at 22°C in both LL and HL conditions (Supplemental Table 2). For Chat-1, Par-4 and Bor-4, HL increased dormancy. However, for Da-0 and Tou-A1-67, it did not show any effect. We next compared germination data immediately after harvesting for all accessions in all conditions (Figure 2). Warm temperature during plant growth reduced dormancy in most of the accessions. Light intensity has a pronounced accession-specific effect on dormancy, when plants are grown at 22°C, from a drastic reduction in dormancy in Mt-0, Col-0 and Ler to no effect in Da-0or Bor-4. The effect of increasing light intensity at 27°C is negligible, except for Tou-A1-67. In general terms, high light intensity reduces dormancy levels, although the effects are accession-specific.

**Figure 2:**
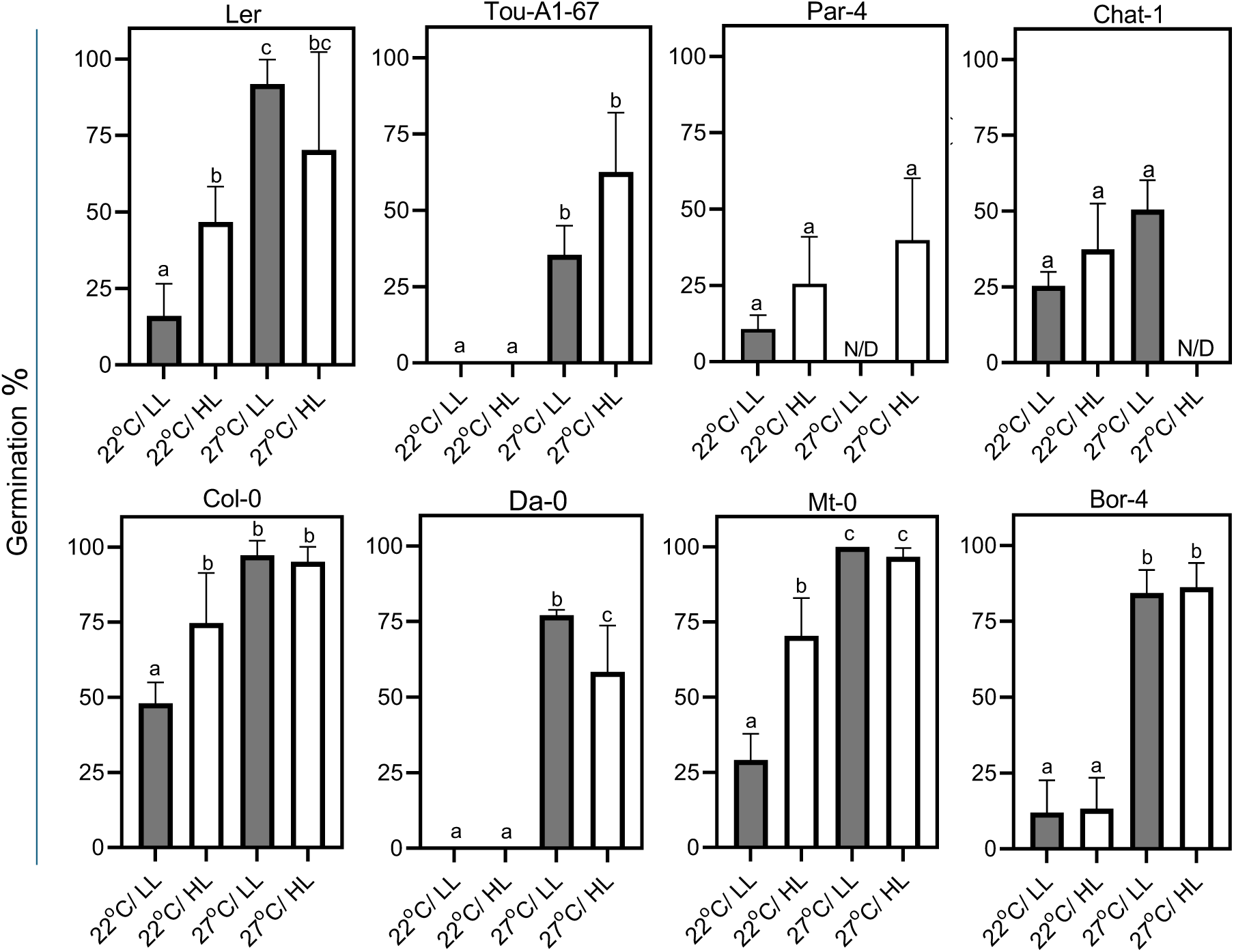
Percentage of germination of *Arabidopsis thaliana* accessions immediately after seed harvest. Data represent mean ± SD for three biological replicates. Significant differences were assessed using a one-way ANOVA followed by a Tukey test (P<0.05). N/D: non-determined due to a lack of enough seeds. LL: low-light conditions. HL: high-light conditions. SD: standard deviation.

### Seed dormancy and longevity relationship is condition-specific

To explore potential associations between dormancy and longevity, we analysed the two traits jointly through correlation analysis. When data for all accessions and all conditions were used, a moderate anticorrelation was found (R^2^=0.33, Supplementary Figure 1), supporting the hypothesis of a negative regulation between both traits (Nguyen et al. 2012, He et al. 2014). A condition-specific analysis showed that this anticorrelation is observed on seeds from plants grown at 22°C under HL conditions (R^2^=0.69) but it’s lost when plants have been grown at 27°C, indicating a strong temperature-specific component in the relationship between both traits (Figure 3). These findings indicate that the correlation between seed dormancy and longevity is dependent on the environmental conditions in which seeds have developed.

**Figure 3:**
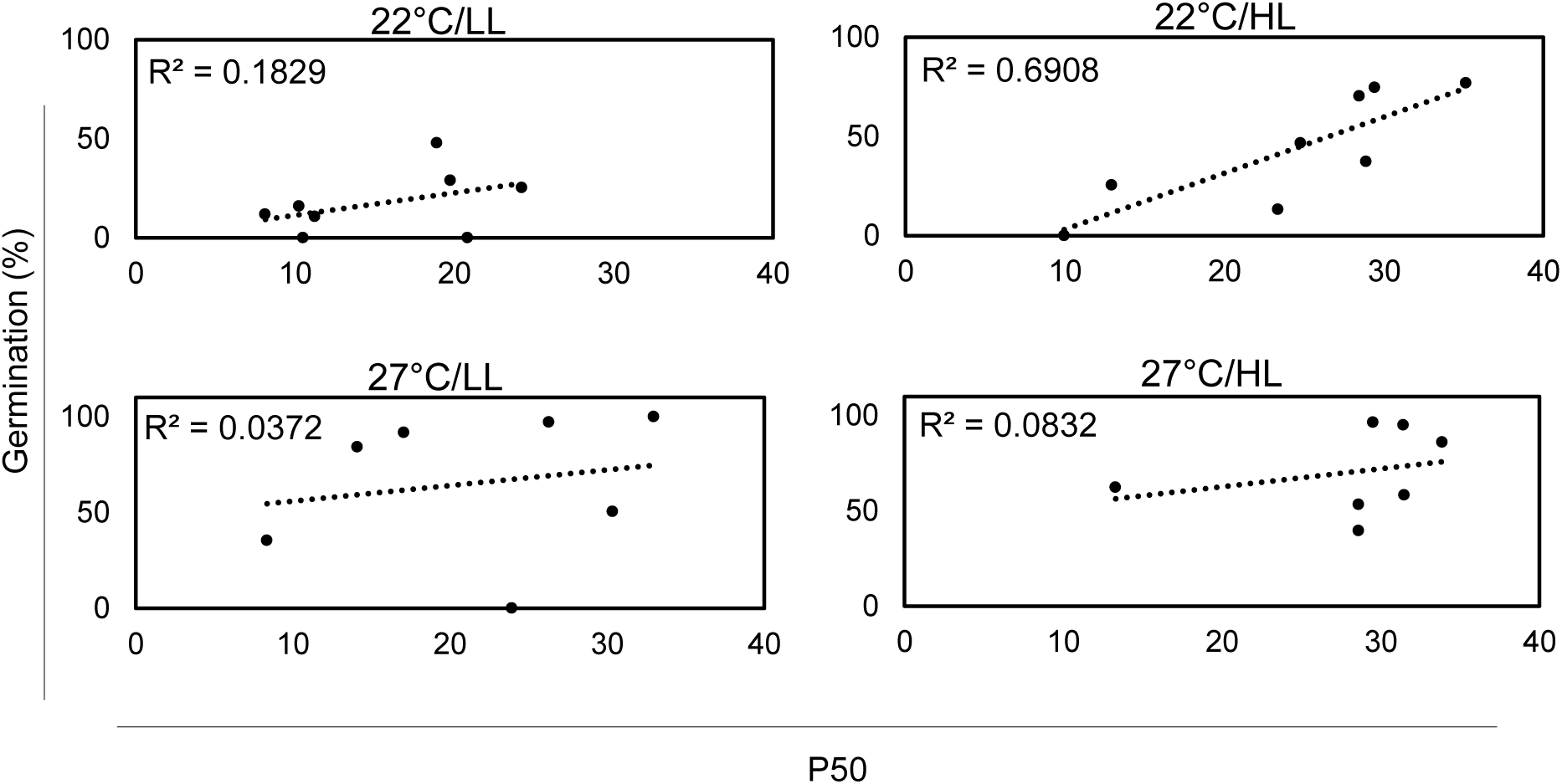
Correlations between P50 values and seed germination (percentage) after harvesting. LL: low-light conditions. HL: high-light conditions.

### Transcriptome analysis of Bor-4 dry seeds reveals a complex reprogramming of stored RNA composition by environmental cues

More than 12,000 different types of stored mRNAs are present in dry seeds of Arabidopsis (Nakabayashi et al. 2005). Differences in the composition of those mRNAs likely influence seed quality parameters (Niñoles et al. 2022). We wanted to investigate how dry seed transcriptomes were modulated by combined environmental signals. We selected Bor-4 for this analysis given its highest plasticity observed for seed longevity (Figure 1). Principal component analysis indicated that both light intensity and temperature modulate Bor-4 dry seed transcriptome, with the effect of temperature being stronger, as evidenced by the separation of the samples in the X-axis (PC1, Principal component 1), which explains 75% of the variability (Figure 4a).

**Figure 4:**
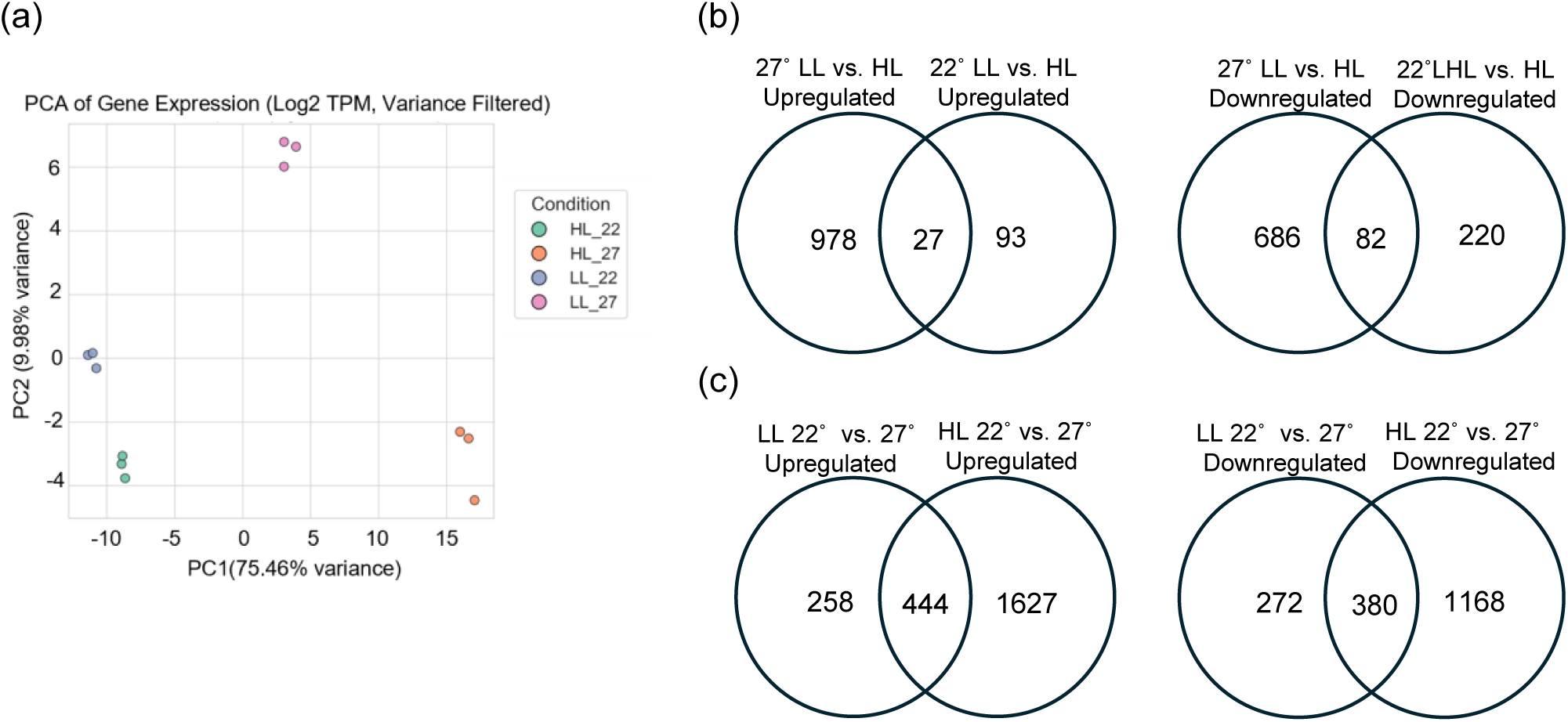
(a) Principal component analysis of transcriptome data of Bor-4 dry seeds grown under four temperature and light conditions (22°C/LL, 22°C/HL, 27°C/LL, 27°C/HL). For every condition, average normalized read count number was employed. The computation includes 75% of variable genes, defined as those with a standard deviation of normalized count data higher than 500 across all four situations. Venn diagram showing upregulated and downregulated genes by light (b) and (c) by temperature in Bor-4 dry seeds grown under four temperature and light conditions (22°C/LL, 22°C/HL, 27°C/LL, 27°C/HL).

When seeds were developed at 22°C, higher light intensity altered the global transcriptome (422 differentially expressed genes, DEGs). However, when developed at 27°C, the number increases dramatically up to 1773 (Supplemental Table 3). A high percentage of these light-dependent changes are temperature-specific (Figure 4b). For example, 89% of the down-regulated, and 97% of the up-regulated genes inducible by high light when seeds are developed at 27°C did not present altered expression when they do at 22°C. Similarly, 72% of the down-regulated, and 77% of the up-regulated genes altered by high light when seed developed at 22°C did not present altered expression when they do at 27°C. A similar trend is observed for temperature-dependent changes. The number of genes with altered expression in dry seeds doubles when seeds are developed at 27°C (Figure 5c), and a high percentage of these temperature-dependent changes are light intensity-specific. 75% of the down-regulated, and 71% of the up-regulated genes altered by high temperature when seeds developed at HL intensity do not present altered expression when they did at LL. Similarly, 58% of the down-regulated, and 35% of the up-regulated genes altered by high temperature when seeds developed at LL do not present altered expression when they did at HL intensity. This indicates that 1) the dry seed transcriptome is very sensitive to light intensity and temperature during seed development, 2) the transcriptional response to high light is potentiated by elevated temperature, and similarly, the response to high temperature is intensified by high light intensity.

**Figure 5:**
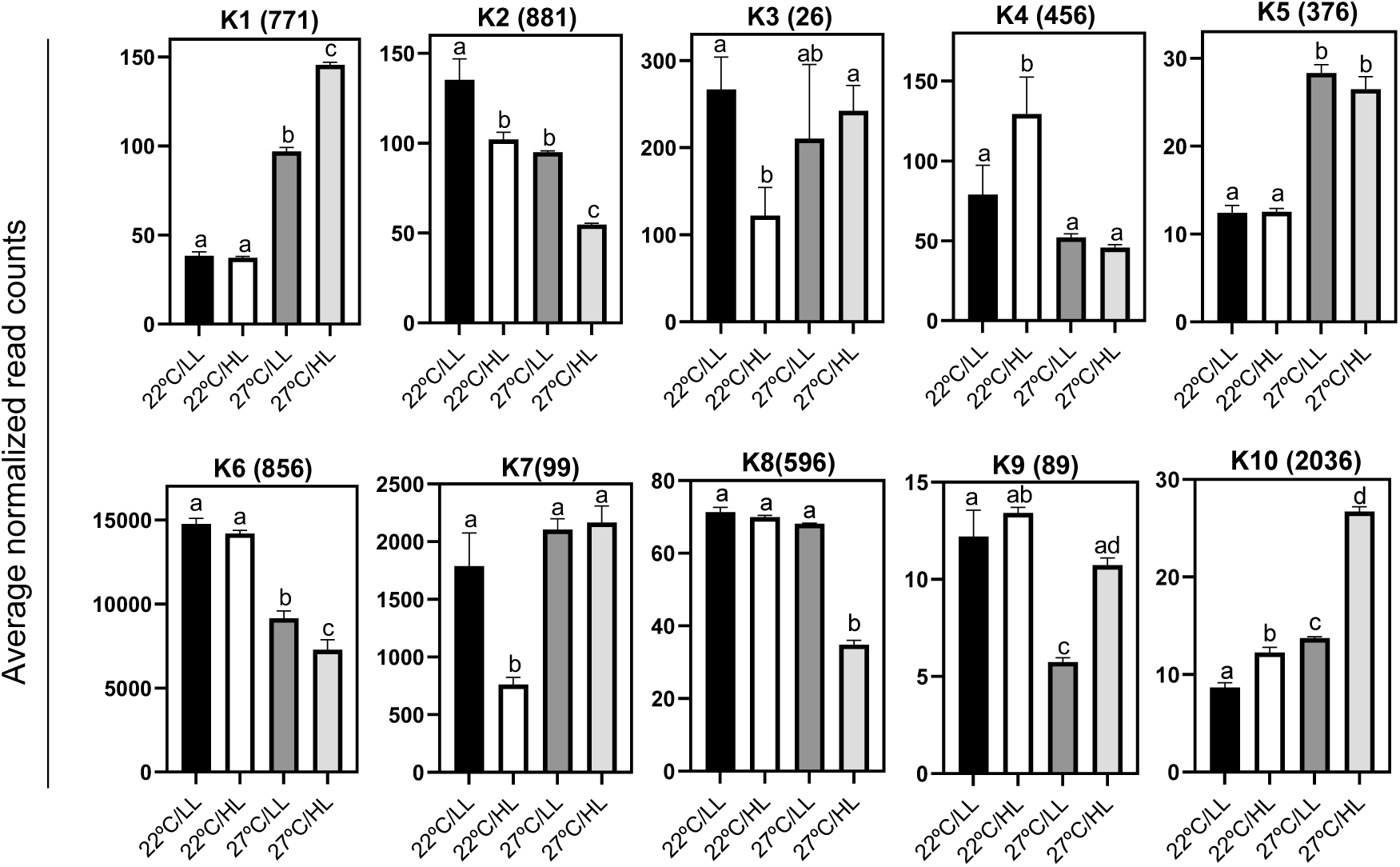
K means clustering of differentially expressed genes of Bor-4 dry seeds grown under four temperature and light conditions (22°C/LL, 22°C/HL, 27°C/LL, 27°C/HL). Number of genes per cluster is in brackets. Y-axis: averaged normalized read counts. Bars represent mean ± SD. Statistical significance was determined by one-way ANOVA followed by a Tukey test, with letters indicating significant differences (P<0.05).

To investigate individual and combined effects, we clustered significantly responding transcripts from the two single signals and the combined treatments and analysed their expression profiles (Figure5, Supplemental Table 4). Responses were classified as prioritized when the response was signal-specific, and as conditional when response to a single signal was dependent on the state of the second one. Combined effects were studied using 22°C/LL seeds as basal conditions and defined when the response to combined signals (HL and 27°C) was different from that of individual ones. Combined effects were further classified as additive (when expression changes by a single signal were potentiated when the two signals were together), cancelled (when the single signal effect was neutralized or attenuated when the two signals were together) or combinatorial (when the response when both signals were applied was different to that to the single ones). 79% of the DEGs presented a combined effect (Supplemental Table 5). Clusters K8 and K10 (55% of them) presented a combinatorial effect. In K8, a decrease in expression was observed only when both signals were together. These clusters include genes related to several responses to abiotic stresses. K10 is the most populated cluster, and includes genes induced only when both signals were together. This cluster includes genes participating in heat stress and oxidative stress responses, photosynthesis (dark reaction) or starch degradation. Clusters K1 and K2 present additive effects (34%) and include temperature-specific genes. K1 includes genes which are induced by high temperature, involved in translation and photosynthesis categories. K2 includes genes repressed by high temperature, involved mainly in stress-related categories. Finally, clusters K3, K4, K7 and K9 present cancelled effects, with K4 being the most populated one. In this cluster, the increase in expression observed under high light conditions at 22°C was no longer observed when the combined signals were applied. This cluster contains genes related to chromatin remodelling (Figure 5; Supplemental Table 6). Overall, a high level of regulatory complexity exists when combined effects are sensed by developing seeds, likely impacting their different future performance. DOG1 expression levels correlated well with dormancy data. A dramatic decrease in DOG1 transcripts was observed in seeds developed at 27°C (the difference more pronounced between 22°C/LL and 27°C/HL seeds), (Supplemental Figure 3a), confirming the role of this key regulator of dormancy in environmental modulation.

### Seeds developed at high light intensity and high temperature are primed with antioxidant defences

We next investigated molecular processes putatively responsible for the extended longevity phenotype observed for Bor-4 seeds. Increments in longevity were found when light intensity was increased but also when temperature was increased at HL or when both signals were applied simultaneously (Figure 1). Among the top 20 enriched categories in the down-regulated genes by HL were found responses to abiotic stresses such as cold, water deprivation or hypoxia. This response was orchestrated independently of the temperature at which seeds were developed, and also when temperature was increased at HL and when both signals were combined (Figure 6a, b, Supplemental Tables 7 and 8). Among the up-regulated genes, however, differences were found dependent on the parental temperature. On seeds developed at 22°C, enrichment was found in categories related to RNA silencing and regulatory small RNAs. On seed developed a 27°C, however, overrepresented categories include responses to heat and to oxidative stress, and cellular respiration, among others (Figure 6c, d). When temperature was increased at HL conditions or both signals were combined, a higher number of significant categories was overrepresented, among them responses to heat, oxidative stress and cellular respiration (Supplemental Table 7 and 8).

**Figure 6:**
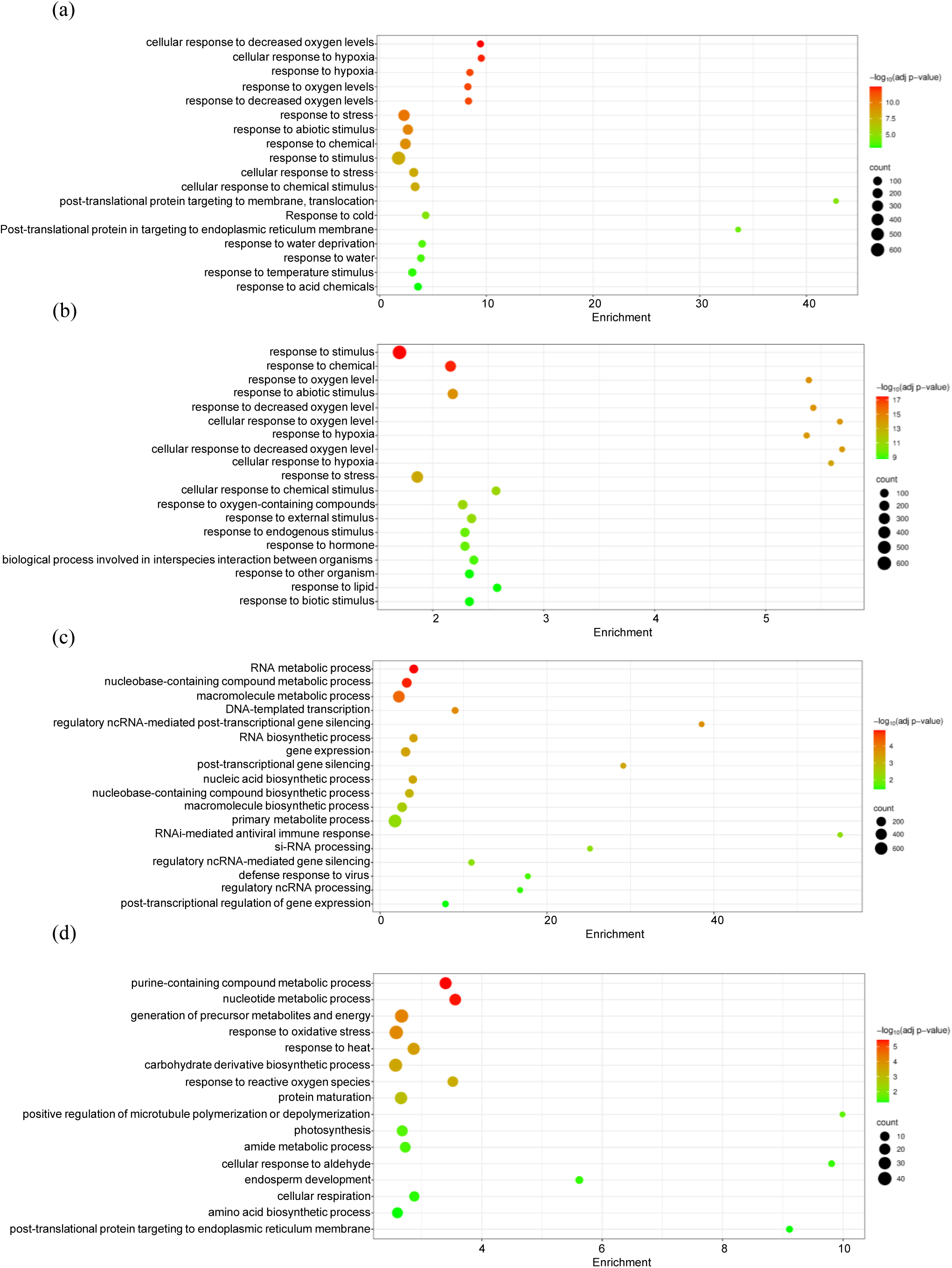
Bubble plots for enriched Gene Ontology (GO) biological processes in (a) genes downregulated by HL at 22°C, (b) genes downregulated by HL at 27°C, (c) genes upregulated by HL at 22°C, and (d) genes upregulated by HL at 27°C. Terms were pruned by REVIGO frequency (terms with frequency > 10% were removed) and ranked by dispensability. Counts, number of genes in a given category. −log P (P-value in log scale after false discovery rate correction).

Given that one of the major threats to longevity is oxidative stress, caused by the accumulation of reactive oxygen species (ROS), damaging cellular components, we focused and extended further the analysis of this response. Cellular respiration category was found enriched in seeds developed under HL, high temperature and combined 27°C/HL signals (Supplemental Tables 7 and 8). We validated this observation using three mitochondrial genes: the NADH dehydrogenase *ATMG00510* gene, the cytochrome C oxidase *ATMG01360* gene, and the *ATMG00640* ATP-synthase subunit gene (Figure 7a). As observed, this mechanism may not be altered in the other three accessions analysed. To further investigate whether higher levels of transcripts could translate into higher mitochondrial activity in these seeds, we measured the levels of the Krebs cycle intermediate citric, malic and fumaric acids in dry and 2h embedded seeds developed at 22°C/HL and 27°C/HL, before and after ageing. As shown in Figure 7b, Krebs cycle intermediates levels are lower in aged seeds developed at 27°C/HL. This trend is also observed when aged seeds are embedded, and the germination process starts, suggesting that mitochondria acclimated at 27°C/HL are more active at respiration (Dellero et al. 2023). Cellular respiration is vital for energy production but inevitably generates ROS as a byproduct (Hernansanz-Agustín and Enríquez 2021), leading to oxidative stress. We assayed oxidative stress in mitochondria of isolated embryos from aged seeds developed under both conditions, but we could not find any significant difference between them (Supplemental Figure 2), suggesting that differences in gene expression in mitochondrial genes does not translate into more oxidative stress on acclimated seeds.

**Figure 7:**
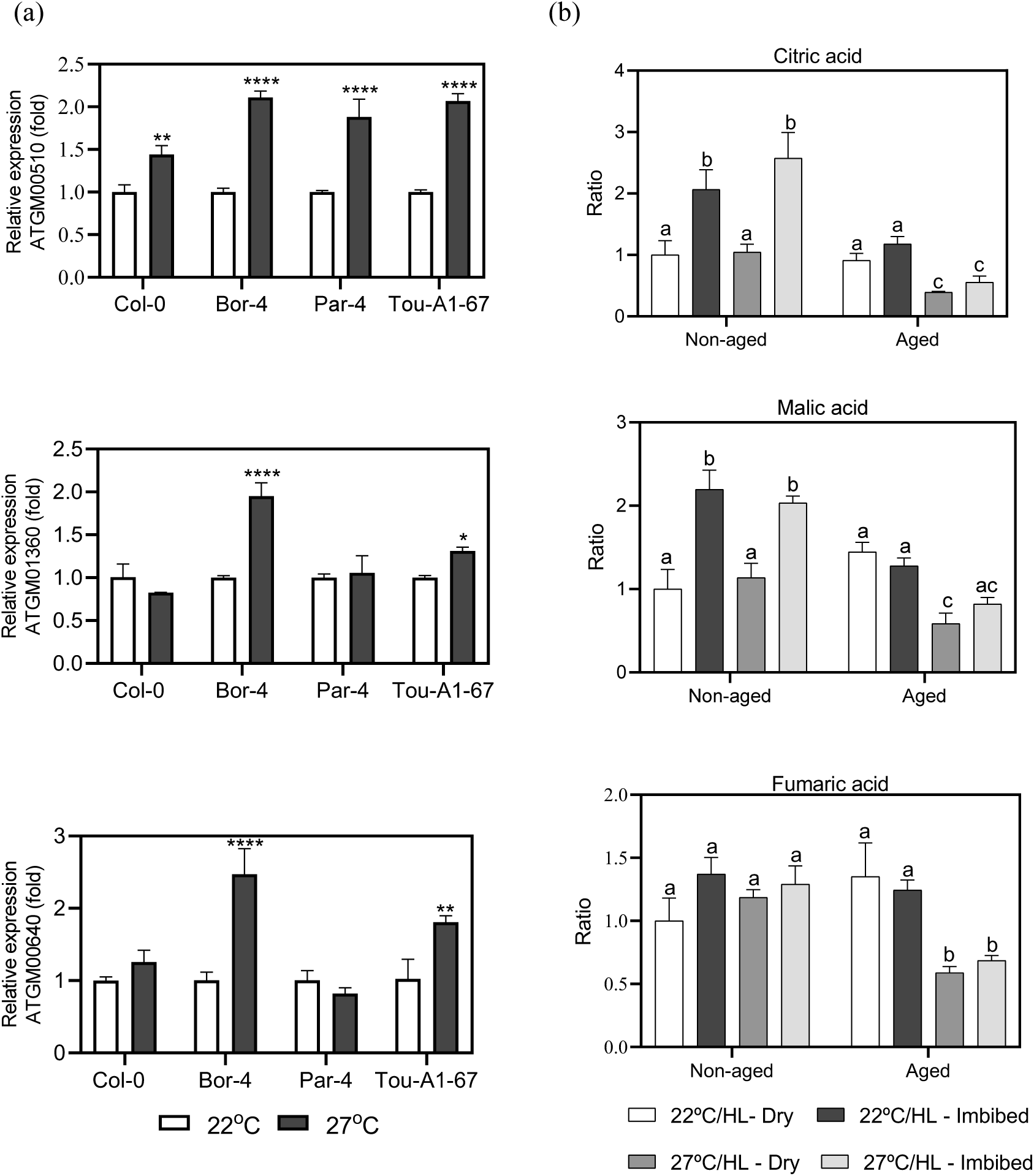
(a) Relative gene expression for NADH dehydrogenase ATMG00510, cytochrome C oxidase ATMG01360, and ATP-synthase subunit ATMG00640 genes. Statistical significance (Student t-test) was determined comparing 22°C/HL and 27°C/HL data within each genotype. Expression at 22°C/HL was normalized to 1 for the three genes. p < 0.01 (**), p < 0.0001 (****), (b) Citric, malic and fumaric acid levels in non-aged and aged dry and imbibed seeds under 22°C/HL or 27°C/HL conditions. Statistical significance was determined using two-way ANOVA followed by a Tukey test, with letters indicating significant differences (p<0.05). Metabolite levels in 22°C/HL dry seeds was normalized to 1. All data represent mean ± SD of three biological replicates.

The scavenging hydrogen peroxide *CAT2* and *CAT3* catalase genes exhibited a dramatic increase in expression when both signals were combined. Similarly, we found increased expression of genes involved in H2O2 detoxification (Supplemental Figure 3b, c). This behaviour was validated for the *CAT3* and for the dehydroascorbate reductase *DHAR1* genes by qRT-PCR (Figure 8a, b). Significant increases were also found in other accessions. Glyoxalases play a major role in methylglyoxal detoxification, a molecule produced as a by-product of lipid peroxidation, leading to oxidative damage of cellular components (Jain et al. 2018). We found several glyoxalase genes more expressed in dry seeds developed at 27°C/HL (Supplemental Figure 3d). This was validated for two genes (*ATGLYI2* and *ATGLX*) also in other accessions (Figure 8c). These results suggest that the activation of antioxidant responses by high light intensity and high temperature is a general mechanism of acclimatation, which could explain the extended longevity phenotype of these seeds.

**Figure 8:**
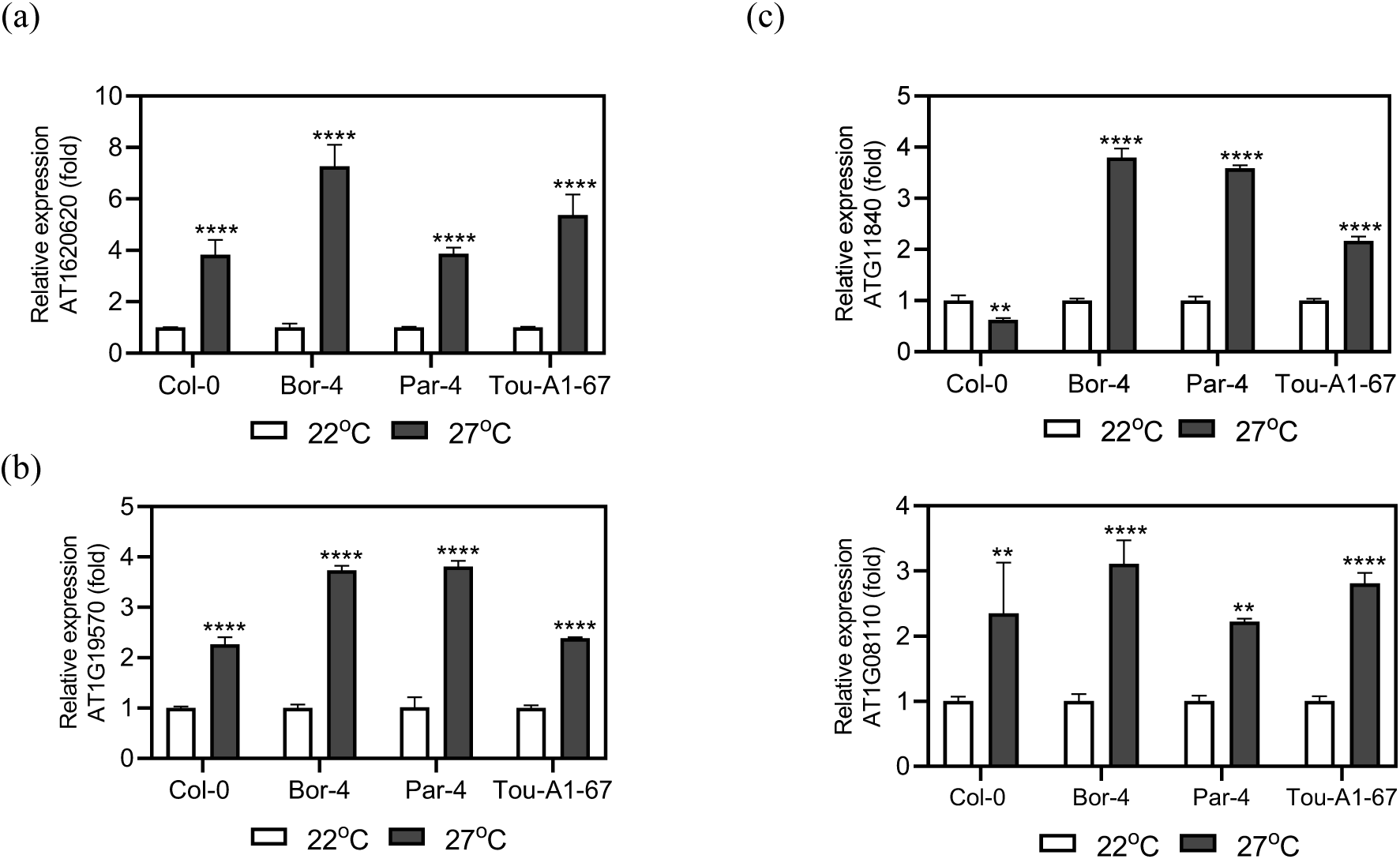
Gene expression by qRT-PCR of (a) catalase 3 CAT3 gene (AT1G20620), (b) dehydroascorbate reductase DHAR1 gene (AT1G119570), (c) glyoxalase genes including ATGLYI2 (AT1G11840) and (d) ATGLX (AT1G08110) in dry seeds developed at 22°C/HL or 27°C/HL. Data represent mean ± SD of three biological replicates. Statistical significance was determined using Student’s t-test comparing 22°C/HL and 27°C/HL conditions within each genotype. p < 0.01 (**), p < 0.0001 (****). Expression at 22°C/HL was normalized to 1 for all genes.

### Raffinose levels increase during ageing in seeds developed at high light and high temperature

Raffinose family oligosaccharides (RFOs) play a significant role in enhancing seed longevity (Li et al. 2017, Salvi et al. 2022), acting as osmoprotectants and stabilizers and protecting against oxidative damage. Their biosynthesis is initiated by the conversion of Myo-inositol into galactinol by means of galactinol synthases enzymes. The galactinol further acts as a galactosyl moiety donor for successive RFO members like raffinose and stachyose, by means of raffinose and stachyose synthases, respectively (Salvi et al. 2022). No clear pattern was observed in the expression levels of the different isoforms of these enzymes in our transcriptome analysis, although induction was observed for galactinol synthase 1 and 5 (*AT2G47180* and *AT5G23790*, Supplemental Table 2). To have a clearer picture of how light intensity and temperature modulate raffinose metabolism on seeds, we measure galactinol and raffinose levels in dry and 2-h embedded seeds developed at 22°C/HL and at 27°C/HL, before and after ageing. Increased light intensity and temperature during seed development have no effect on the concentration of galactinol and raffinose of dry seeds (Figure 9). After ageing, the levels of galactinol decreased and the levels of raffinose increased. Interestingly, seeds developed at 27°C/HL accumulated more raffinose than those developed at 22°C/HL after the ageing treatment. This suggests that acclimated seeds at 27°C/HL have a higher potential to deploy RFOs and to safeguard cellular components under situations generating oxidative damage.

**Figure 9:**
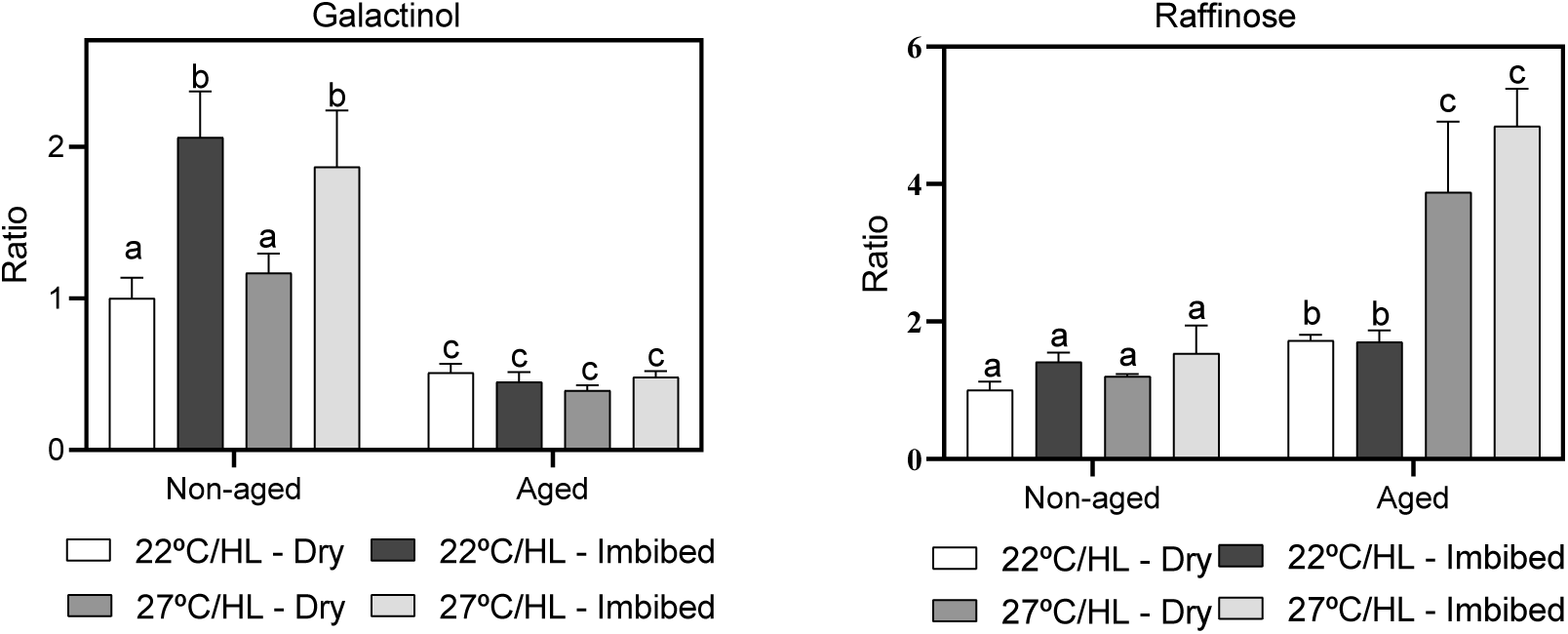
Galactinol and raffinose relative levels in non-aged and aged dry and imbibed seeds under 22°C/HL or 27°C/HL conditions. Data represent mean ± SD of three biological replicates. Statistical differences between non-aged and aged within each treatment were determined using a two-way ANOVA followed by a Tukey test, with letters indicating significant differences (P<0.05). Metabolite levels were normalized to 1 in 22°C/HL dry seeds.

## Discussion

Climate change typically entails the simultaneous occurrence of multiple factors, such as heat and altered light conditions. Warming alters the diurnal asymmetry in cloud coverage, increasing the daytime solar radiation while enhancing or diminishing net temperature response depending on latitude and diurnal cycle strength (Brient and Bony 2013, Luo et al. 2024). This complex interplay of fluctuations in the environment will likely affect the future seeds in a combinatorial way. Acclimation, defined as the ability to regulate plant physiology in response to mild environmental changes during prolonged periods, causes less damage than short-term stress, and triggers adjustments for long-term optimization of resources (Klupczyńska and Pawłowski 2021). Our experimental design using contrasting parental environments, combining light intensities and temperatures within a physiological range, has enabled us to gain a comprehensive view of the impact of these two signals on seed acclimation. In addition, analysing eight distinct natural accessions allowed us to estimate the weight of the genetic background on the variation of two key seed characteristics, dormancy and longevity, without compromising conclusions to the study to just one genotype.

Temperature can impact longevity positively or negatively in a genetic-dependent manner. Warm temperatures are detrimental to longevity in *Wahlenbergia* or *Medicago (Kochanek et al. 2009, Righetti et al. 2015)* but beneficial for *Plantago* (Kochanek et al. 2011). In Arabidopsis, contradictory results have been reported so far. He et al. 2014)observed that 27°C parental environment generated seeds with increased longevity. Malabarba et al. 2021, however, found a detrimental effect of the same parental temperature on seeds of the same accession (Col-0). In this study, seeds from different accessions (including Col-0) developed at a combination of high temperature (27°C) and high light intensity exhibited a common trend of increased longevity. Altogether, these observations suggest that the boundary between positive and negative effects of temperature on longevity is very narrow and likely depends on other unknown experimental variables. This is not unexpected, given the polygenic nature of this trait (Niñoles et al. 2022). Temperature likely has both detrimental and beneficial effects on the seed. Malabarba et al. 2021 reported increases in suberin biosynthesis and heat-shock proteins in seeds developed at 27°C, known to be beneficial for longevity (Kaur et al. 2015, Almoguera et al. 2020, Renard et al. 2020, Niñoles et al. 2022, Wang et al. 2024) but also decreases in ribosome biogenesis genes, which is detrimental (Wang et al. 2023, Lyu et al. 2025). Similarly, high temperature stimulates reactive-oxygen species production, which will cause oxidative stress and would accelerate deterioration (Soengas et al. 2018), but it also activates the expression of antioxidant defences (Liu et al. 2021). The equilibrium between beneficial and adverse effects elicited by mild increases in temperature seems a critical factor in determining the overall physiological outcome for seed longevity. A positive effect of temperature on seed longevity was also found by (Mondoni et al. 2011). *Silene vulgaris* seeds from lowland locations exhibited higher longevity than those from alpine locations, correlating with higher expression of the *SvHSP17.4* heat-shock protein gene. This example further illustrates the importance of genotype and combined environmental effects in determining the outcome. Despite alpine seeds receiving higher light intensity, which would be beneficial for longevity, at least in Arabidopsis ((He et al. 2014); Figure1), the higher temperature of lowland seeds could have a predominant impact on longevity for this species. Heat-shock proteins (HSP) have a clear positive effect on seed longevity (Prieto-Dapena et al. 2006, Wozny et al. 2018, Niñoles et al. 2022), suggesting that the higher levels of HSP in our acclimated seeds (Supplemental Table 3) will potentiate the increased resistance to seed deterioration observed in these seeds. Genotype plasticity has been reported multiple times for dormancy, which is regulated by temperature and light in a genotype-specific manner (Schmuths et al. 2006, Vidigal et al. 2016, Kerdaffrec and Nordborg 2017, Huang et al. 2019, Martínez-Berdeja et al. 2020, Iwasaki et al. 2022, Bonnot et al. 2023). For example, although transcript and protein abundance of the major dormancy gene determinant *DOG1* generally correlate with seed dormancy, this association is not consistently observed across all accessions (Chiang et al. 2011, Nakabayashi et al. 2012). We found differences in plasticity for dormancy and longevity in the eight accessions assayed, reinforcing a role of the genetic background in the outcome for these two traits. The observation that the most pronounced increases in longevity occurred in accessions classified as “short-lived” points to the involvement of shared protective mechanisms, potentially already present in the “long-lived” accessions but lacking or attenuated in the short-lived ones. The multifactorial nature of seed longevity is also illustrated when levels of expression of the *DOG1* gene in the Bor-4 accession are analysed in the different environments. Lower levels of dormancy correlate well with decreases in *DOG* transcript abundance (Supplemental Figure 3a). However, this does not translate into reduced longevity, as the highest longevity was found in the least dormant 27°C/HL seeds (Figure 1) even though it is known that the *dog1* mutant of Arabidopsis shows reduced longevity (Bentsink et al. 2006). Late maturation genes, including late embryogenesis-related (LEA) and HSP coding-genes, as well as RFO biosynthesis genes are less abundant in dry seeds of the *dog1* mutant, which could explain the reduced longevity genotype (Arroyo-Mosso et al. 2025). Clearly, the higher abundance of transcripts related to ribosome biogenesis and translation categories when *DOG1* is not expressed (Dekkers et al. 2016) is not enough to generate a compensatory effect. Altogether, this indicates that the increased longevity phenotype observed in 27°C/HL seeds is independent or just partially dependent on *DOG1* expression.

The challenge of assigning a phenotype to a particular pathway is highlighted when the combinatorial impact of light and temperature cues is examined (Huang et al. 2019). When Bor-4 seeds developed in the four environmental regimes were compared, 79% of the DEGs presented combined effects. A substantial number of stress-related genes showed decreased expression in 27°C/HL seeds, suggesting that despite the detrimental effects of high light and high temperature, the seeds have been acclimated (Karim and Johnson 2021). Under prolonged light stress, this decrease in stress-related gene expression is typically associated with decreased investments in antioxidant systems (Gollan and Aro 2020, Wang et al. 2020), as the oxidative damage generated by the stress is mitigated by the increase in the ability to adjust the composition of their photosynthetic apparatuses through acclimation (Walters 2005, Herrmann et al. 2019). In our seeds developed at combined 27°C/HL, the lower expression of stress-related genes in clusters K8 and K2 suggests that these seeds are acclimated to these environmental conditions, further inferred by the up-regulation of photosynthetic genes in K1. Interestingly, this is here accompanied by a higher abundance of transcripts of antioxidant defences and translation in K10 and K1 clusters, that exhibit combinatorial and additive behaviours. A distinct ROS signature when stresses are combined has been previously reported (Choudhury et al. 2017).

Seed aging is correlated with a decline in the cellular antioxidant potential (Kranner et al. 2006). Therefore, the high levels of oxidative defences provided by the combined environmental signals could be crucial to generate a highly protective state in these seeds (Bailly 2004, Kibinza et al. 2011, Renard et al. 2020, Zhang et al. 2021). We found higher levels of transcripts involved in cellular respiration in 27°C/HL seeds, which could eventually translate into higher mitochondrial metabolic activity, but we did not observe more ROS in these mitochondria (Figure 7a, Supplemental Figure 2), reinforcing the hypothesis of higher antioxidant protection in these seeds. Antioxidant responses are diverse, encompassing a wide range of cellular pathways and mechanisms to combat oxidative stress. Antioxidant enzymes include superoxide dismutase, ascorbate peroxidase, catalase, dehydroascorbate reductase or monodehydroascorbate reductase (Møller et al. 2007). Catalase (Goel and Sheoran 2003, Lehner et al. 2008, Yin et al. 2014, Sun et al. 2022, Lv et al. 2024, Sun et al. 2024) and dehydroascorbate reductase and monodehydroascorbate reductase activity (Cheng et al. 2020, Sun et al. 2022, Lv et al. 2024, Sun et al. 2024) decline during seed aging, and seeds form dehydroascorbate reductase mutant lines exhibit reduced longevity (Renard et al. 2020). We also found higher expression of glyoxalase (*GLX*) genes in seeds exhibiting enhanced longevity (Supplemental Figure 3). Methylglyoxal (MG) is a potential cytotoxin disrupting cellular functions including the peroxidation of lipids, the oxidation of proteins or the oxidation of fatty acids (Li 2016). The GLX system, the primary route for detoxification of MG, is also reported to contribute to seed longevity. GLX activity increased during ageing of oat seeds (Sun et al. 2022) and seeds from rice transgenic lines with overexpression and disruption of the OsGLYI3 gene exhibited higher and lower germination after accelerated ageing (Liu et al. 2022). Finally, RFOs have been also correlated with seed longevity (de Souza Vidigal et al. 2016, He et al. 2016, Salvi et al. 2022), likely acting as antioxidants to counteract the accumulation of ROS (Nishizawa-Yokoi et al. 2008). Overexpression of the maize alkaline α-galactosidase 1 ZmAGA1 protein in Arabidopsis decreases both RFOs and galactinol contents and reduces seed aging tolerance (Zhang et al. 2021). RFO accumulation was affected in *Medicago abi5* mutants, concomitant with the decrease in longevity, reinforcing the genetic link between these two traits (Zinsmeister et al. 2016). The levels of galactinol decreased and the levels of raffinose increased during ageing in our seeds. Interestingly, seeds developed at 27°C/HL accumulated more raffinose than those developed at 22°C/HL after ageing, which could confer an additional advantage to these seeds upon ageing-induced oxidative damage. Overall, combined high light intensity and high temperature may endow seeds with a higher antioxidant potential in the form of stored mRNAs, which makes them better equipped to withstand aging. These findings reinforce the crucial role of antioxidant defences in seed longevity and position them as promising targets for future biotechnological approaches aimed at improving seed viability.

## Data availability

The datasets generated for this study can be found in the GEO database under the accession number GSE277583.

## Author contributions

SMA, DV, EY, PM, JF, ST: Investigation– review & editing., RG, CG: Conceptualization, Formal analysis, Writing – review & editing, JG: Conceptualization, Formal analysis, Validation, Writing – original draft, Writing

## References

Albanese, P., M. Manfredi, A. Meneghesso, E. Marengo, G. Saracco, J. Barber, T. Morosinotto and C. Pagliano (2016). Dynamic reorganization of photosystem II supercomplexes in response to variations in light intensities. Biochimica et Biophysica Acta (BBA)-Bioenergetics, 1857(10), 1651–1660.

Almoguera, C., P. Prieto-Dapena, R. Carranco, J. L. Ruiz and J. Jordano (2020). Heat stress factors expressed during seed maturation differentially regulate seed longevity and seedling greening. Plants, 9(3), 335.

Andalo, C., S. Mazer, B. Godelle and N. Machon (1999). Parental environmental effects on life history traits in Arabidopsis thaliana (Brassicaceae). The New Phytologist, 142(2), 173–184.

Andrews, S. 2010. FastQC: A Quality Control Tool for High Throughput Sequence Data http://www.bioinformatics.babraham.ac.uk/projects/fastqc/

Arroyo-Mosso, I. A., H. N. Diaz-Ardila, A. Garciarrubio, U. Kumara, D. F. Rendón-Luna, T. B. Nava-Ramírez, T. C. Boothby, J. L. Reyes and A. A. Covarrubias (2025). A Group 6 LEA Protein Plays Key Roles in Tolerance to Water Deficit, and in Maintaining the Glassy State and Longevity of Seeds. Plant, Cell & Environment.

Bailly, C. (2004). Active oxygen species and antioxidants in seed biology. Seed science research, 14(2), 93–107.

Balfagón, D., S. Sengupta, A. Gómez-Cadenas, F. B. Fritschi, R. K. Azad, R. Mittler and S. I. Zandalinas (2019). Jasmonic acid is required for plant acclimation to a combination of high light and heat stress. Plant physiology, 181(4), 1668–1682.

Balfagón, D., S. I. Zandalinas, T. dos Reis de Oliveira, C. Santa-Catarina and A. Gómez-Cadenas (2022). Reduction of heat stress pressure and activation of photosystem II repairing system are crucial for citrus tolerance to multiple abiotic stress combination. Physiologia plantarum, 174(6), e13809.

Ballottari, M., L. Dall’Osto, T. Morosinotto and R. Bassi (2007). Contrasting behavior of higher plant photosystem I and II antenna systems during acclimation. Journal of Biological Chemistry, 282(12), 8947–8958.

Bentsink, L., J. Jowett, C. J. Hanhart and M. Koornneef (2006). Cloning of DOG1, a quantitative trait locus controlling seed dormancy in Arabidopsis. Proceedings of the national academy of sciences, 103(45), 17042–17047.

Bonnot, T., I. Somayanda, S. K. Jagadish and D. H. Nagel (2023). Time of day and genotype sensitivity adjust molecular responses to temperature stress in sorghum. The Plant Journal, 116(4), 1081–1096.

Brient, F. and S. Bony (2013). Interpretation of the positive low-cloud feedback predicted by a climate model under global warming. Climate Dynamics, 40(9), 2415–2431.

Casal, J. J. and S. Balasubramanian (2019). Thermomorphogenesis. Annual review of plant biology, 70(1), 321–346.

Chen, M., D. R. MacGregor, A. Dave, H. Florance, K. Moore, K. Paszkiewicz, N. Smirnoff, I. A. Graham and S. Penfield (2014). Maternal temperature history activates Flowering Locus T in fruits to control progeny dormancy according to time of year. Proceedings of the National Academy of Sciences, 111(52), 18787–18792.

Cheng, H., X. Ma, S. Jia, M. Li and P. Mao (2020). Transcriptomic analysis reveals the changes of energy production and AsA-GSH cycle in oat embryos during seed ageing. Plant Physiology and Biochemistry, 153, 40–52.

Chiang, G. C., M. Bartsch, D. Barua, K. Nakabayashi, M. Debieu, I. Kronholm, M. Koornneef, W. J. Soppe, K. Donohue and J. De Meaux (2011). DOG1 expression is predicted by the seed-maturation environment and contributes to geographical variation in germination in Arabidopsis thaliana. Molecular ecology, 20(16), 3336–3349.

Choudhury, F. K., R. M. Rivero, E. Blumwald and R. Mittler (2017). Reactive oxygen species, abiotic stress and stress combination. The Plant Journal, 90(5), 856–867.

Clerkx, E. J., H. B.-D. Vries, G. J. Ruys, S. P. Groot and M. Koornneef (2003). Characterization of green seed, an enhancer of abi3-1 in Arabidopsis that affects seed longevity. Plant physiology, 132(2), 1077–1084.

Crawford, A. J., D. H. McLachlan, A. M. Hetherington and K. A. Franklin (2012). High temperature exposure increases plant cooling capacity. Current Biology, 22(10), R396–R397.

Czechowski, T., M. Stitt, T. Altmann, M. K. Udvardi and W.-R. Scheible (2005). Genome-wide identification and testing of superior reference genes for transcript normalization in Arabidopsis. Plant physiology, 139(1), 5–17.

de Souza Vidigal, D., L. Willems, J. van Arkel, B. J. Dekkers, H. W. Hilhorst and L. Bentsink (2016). Galactinol as marker for seed longevity. Plant Science, 246, 112–118.

Debeaujon, I., K. M. Leon-Kloosterziel and M. Koornneef (2000). Influence of the testa on seed dormancy, germination, and longevity in Arabidopsis. Plant physiology, 122(2), 403–414.

Dekkers, B. J., H. He, J. Hanson, L. A. Willems, D. C. Jamar, G. Cueff, L. Rajjou, H. W. Hilhorst and L. Bentsink (2016). The Arabidopsis DELAY OF GERMINATION 1 gene affects ABSCISIC ACID INSENSITIVE 5 (ABI 5) expression and genetically interacts with ABI 3 during Arabidopsis seed development. The Plant Journal, 85(4), 451–465.

Dellero, Y., S. Berardocco, C. Berges, O. Filangi and A. Bouchereau (2023). Validation of carbon isotopologue distribution measurements by GC-MS and application to 13C-metabolic flux analysis of the tricarboxylic acid cycle in Brassica napus leaves. Frontiers in plant science, 13, 885051.

Dobin A, Davis CA, Schlesinger F, Drenkow J, Zaleski C, Jha S, Batut P, Chaisson M, Gingeras TR. STAR: ultrafast universal RNA-seq aligner. Bioinformatics. 2013 Jan 1;29(1):15–21. doi: 10.1093/bioinformatics/bts635

Goel, A. and I. Sheoran (2003). Lipid peroxidation and peroxide-scavenging enzymes in cotton seeds under natural ageing. Biologia plantarum, 46(3), 429–434.

Gollan, P. J. and E.-M. Aro (2020). Photosynthetic signalling during high light stress and: targets and dynamics. Philosophical Transactions of the Royal Society B, 375(1801), 20190406.

He, H., D. de Souza Vidigal, L. B. Snoek, S. Schnabel, H. Nijveen, H. Hilhorst and L. Bentsink (2014). Interaction between parental environment and genotype affects plant and seed performance in Arabidopsis. Journal of experimental botany, 65(22), 6603–6615.

He, H., L. A. Willems, A. Batushansky, A. Fait, J. Hanson, H. Nijveen, H. W. Hilhorst and L. Bentsink (2016). Effects of parental temperature and nitrate on seed performance are reflected by partly overlapping genetic and metabolic pathways. Plant and Cell Physiology, 57(3), 473– 487.

Hernansanz-Agustín, P. and J. A. Enríquez (2021). Generation of reactive oxygen species by mitochondria. Antioxidants, 10(3), 415.

Herrmann, H. A., J.-M. Schwartz and G. N. Johnson (2019). Metabolic acclimation—a key to enhancing photosynthesis in changing environments? Journal of experimental botany, 70(12), 3043–3056.

Huang, J., X. Zhao and J. Chory (2019). The Arabidopsis transcriptome responds specifically and dynamically to high light stress. Cell reports, 29(12), 4186–4199. e4183.

Huang, Z., S. Footitt, A. Tang and W. E. Finch-Savage (2018). Predicted global warming scenarios impact on the mother plant to alter seed dormancy and germination behaviour in Arabidopsis. Plant, cell & environment, 41(1), 187–197.

Iwasaki, M., S. Penfield and L. Lopez-Molina (2022). Parental and environmental control of seed dormancy in Arabidopsis thaliana. Annual Review of Plant Biology, 73(1), 355–378.

Jain, M., P. Nagar, A. Sharma, R. Batth, S. Aggarwal, S. Kumari and A. Mustafiz (2018). GLYI and D-LDH play key role in methylglyoxal detoxification and abiotic stress tolerance. Scientific reports, 8(1), 5451.

Kan, Y., X.-R. Mu, J. Gao, H.-X. Lin and Y. Lin (2023). The molecular basis of heat stress responses in plants. Molecular Plant, 16(10), 1612–1634.

Karim, M. F. and G. N. Johnson (2021). Acclimation of photosynthesis to changes in the environment results in decreases of oxidative stress in Arabidopsis thaliana. Frontiers in Plant Science, 12, 683986.

Kaur, H., B. P. Petla, N. U. Kamble, A. Singh, V. Rao, P. Salvi, S. Ghosh and M. Majee (2015). Differentially expressed seed aging responsive heat shock protein OsHSP18. 2 implicates in seed vigor, longevity and improves germination and seedling establishment under abiotic stress. Frontiers in Plant Science, 6, 713.

Kendall, S. L., A. Hellwege, P. Marriot, C. Whalley, I. A. Graham and S. Penfield (2011). Induction of dormancy in Arabidopsis summer annuals requires parallel regulation of DOG1 and hormone metabolism by low temperature and CBF transcription factors. The Plant Cell, 23(7), 2568–2580.

Kerdaffrec, E. and M. Nordborg (2017). The maternal environment interacts with genetic variation in regulating seed dormancy in Swedish Arabidopsis thaliana. PloS one, 12(12), e0190242.

Kibinza, S., J. Bazin, C. Bailly, J. M. Farrant, F. Corbineau and H. El-Maarouf-Bouteau (2011). Catalase is a key enzyme in seed recovery from ageing during priming. Plant science, 181(3), 309–315.

Klupczyńska, E. A. and T. A. Pawłowski (2021). Regulation of seed dormancy and germination mechanisms in a changing environment. International journal of molecular sciences, 22(3), 1357.

Kochanek, J., K. J. Steadman, R. J. Probert and S. W. Adkins (2009). Variation in seed longevity among different populations, species and genera found in collections from wild Australian plants. Australian Journal of Botany, 57(2), 123–131.

Kochanek, J., K. J. Steadman, R. J. Probert and S. W. Adkins (2011). Parental effects modulate seed longevity: exploring parental and offspring phenotypes to elucidate pre-zygotic environmental influences. New Phytologist, 191(1), 223–233.

Koini, M. A., L. Alvey, T. Allen, C. A. Tilley, N. P. Harberd, G. C. Whitelam and K. A. Franklin (2009). High temperature-mediated adaptations in plant architecture require the bHLH transcription factor PIF4. Current biology, 19(5), 408–413.

Kranner, I., S. Birtić, K. M. Anderson and H. W. Pritchard (2006). Glutathione half-cell reduction potential: a universal stress marker and modulator of programmed cell death? Free Radical Biology and Medicine, 40(12), 2155–2165.

Lee, H., K. Calvin, D. Dasgupta, G. Krinner, A. Mukherji, P. Thorne, C. Trisos, J. Romero, P. Aldunce and K. Barrett (2023). Climate change 2023: synthesis report. Contribution of working groups I, II and III to the sixth assessment report of the intergovernmental panel on climate change.

Lehner, A., N. Mamadou, P. Poels, D. Côme, C. Bailly and F. Corbineau (2008). Changes in soluble carbohydrates, lipid peroxidation and antioxidant enzyme activities in the embryo during ageing in wheat grains. Journal of Cereal Science, 47(3), 555–565.

Li, T., Y. Zhang, D. Wang, Y. Liu, L. M. Dirk, J. Goodman, A. B. Downie, J. Wang, G. Wang and T. Zhao (2017). Regulation of seed vigor by manipulation of raffinose family oligosaccharides in maize and Arabidopsis thaliana. Molecular Plant, 10(12), 1540–1555.

Li, Z.-G. (2016). Methylglyoxal and glyoxalase system in plants: old players, new concepts. The Botanical Review, 82(2), 183–203.

Liu, H.-L., Z.-X. Lee, T.-W. Chuang and H.-C. Wu (2021). Effect of heat stress on oxidative damage and antioxidant defense system in white clover (Trifolium repens L.). Planta, 254(5), 103.

Liu, S., W. Liu, J. Lai, Q. Liu, W. Zhang, Z. Chen, J. Gao, S. Song, J. Liu and Y. Xiao (2022). OsGLYI3, a glyoxalase gene expressed in rice seed, contributes to seed longevity and salt stress tolerance. Plant Physiology and Biochemistry, 183, 85–95.

Liu, Y., S. Liu, Y. Jing, J. Li and R. Lin (2025). Light regulates seed dormancy through FHY3-mediated activation of ACC OXIDASE 1 in Arabidopsis. Plant Mol Biol, 13(115).

Love MI, Huber W, Anders S. 2014. “Moderated estimation of fold change and dispersion for RNA-seq data with DESeq2.” Genome Biology, 15, 550. d

Luo, H., J. Quaas and Y. Han (2024). Diurnally asymmetric cloud cover trends amplify greenhouse warming. Science Advances, 10(25), eado5179.

Lv, T., J. Li, L. Zhou, T. Zhou, H. W. Pritchard, C. Ren, J. Chen, J. Yan and J. Pei (2024). Aging-Induced Reduction in Safflower Seed Germination via Impaired Energy Metabolism and Genetic Integrity Is Partially Restored by Sucrose and DA-6 Treatment. Plants, 13(5), 659.

Lyu, Y., W. He, X. Zhou, X. Gao, M. Fan, D. Chen and X. Chen (2025). Clathrin Light Chain 2 is a Substrate of Ubiquitin Ligase ATL5 and Negatively Regulates Seed Longevity in Arabidopsis. Journal of Experimental Botany, eraf231.

MacGregor, D. R., S. L. Kendall, H. Florance, F. Fedi, K. Moore, K. Paszkiewicz, N. Smirnoff and S. Penfield (2015). Seed production temperature regulation of primary dormancy occurs through control of seed coat phenylpropanoid metabolism. New Phytologist, 205(2), 642–652.

Malabarba, J., D. Windels, W. Xu and J. Verdier (2021). Regulation of DNA (de) methylation positively impacts seed germination during seed development under heat stress. Genes, 12(3), 457.

Martin, M. 2011. Cutadapt removes adapter sequences from high-throughput sequencing reads. EMBnet.journal,17, n. 1, pp. 10–12

Martínez-Berdeja, A., M. C. Stitzer, M. A. Taylor, M. Okada, E. Ezcurra, D. E. Runcie and J. Schmitt (2020). Functional variants of DOG1 control seed chilling responses and variation in seasonal life-history strategies in Arabidopsis thaliana. Proceedings of the National Academy of Sciences, 117(5), 2526–2534.

Mittler, R. (2006). Abiotic stress, the field environment and stress combination. Trends in plant science, 11(1), 15–19.

Mizutani, M. and M. M. Kanaoka (2018). Environmental sensing and morphological plasticity in plants. Seminars in Cell & Developmental Biology, Elsevier.

Møller, I. M., P. E. Jensen and A. Hansson (2007). Oxidative modifications to cellular components in plants. Annu. Rev. Plant Biol., 58(1), 459–481.

Mondoni, A., R. J. Probert, G. Rossi, E. Vegini and F. R. Hay (2011). Seeds of alpine plants are short lived: implications for long-term conservation. Annals of botany, 107(1), 171–179.

Muhlenbock, P., M. Szechynska-Hebda, M. Płaszczyca, M. Baudo, A. Mateo, P. M. Mullineaux, J. E. Parker, B. Karpinska and S. Karpiński (2008). Chloroplast signaling and LESION SIMULATING DISEASE1 regulate crosstalk between light acclimation and immunity in Arabidopsis. The Plant Cell, 20(9), 2339–2356.

Nakabayashi, K., M. Bartsch, Y. Xiang, E. Miatton, S. Pellengahr, R. Yano, M. Seo and W. J. Soppe (2012). The time required for dormancy release in Arabidopsis is determined by DELAY OF GERMINATION1 protein levels in freshly harvested seeds. The Plant Cell, 24(7), 2826–2838.

Nakabayashi, K., M. Okamoto, T. Koshiba, Y. Kamiya and E. Nambara (2005). Genome-wide profiling of stored mRNA in Arabidopsis thaliana seed germination: epigenetic and genetic regulation of transcription in seed. The plant journal, 41(5), 697–709.

Nakashima, K., Y. Fujita, N. Kanamori, T. Katagiri, T. Umezawa, S. Kidokoro, K. Maruyama, T. Yoshida, K. Ishiyama and M. Kobayashi (2009). Three Arabidopsis SnRK2 protein kinases, SRK2D/SnRK2. 2, SRK2E/SnRK2. 6/OST1 and SRK2I/SnRK2. 3, involved in ABA signaling are essential for the control of seed development and dormancy. Plant and Cell Physiology, 50(7), 1345–1363.

Nguyen, T.-P., P. Keizer, F. van Eeuwijk, S. Smeekens and L. Bentsink (2012). Natural variation for seed longevity and seed dormancy are negatively correlated in Arabidopsis. Plant physiology, 160(4), 2083–2092.

Niñoles, R., D. Planes, P. Arjona, C. Ruiz-Pastor, R. Chazarra, J. Renard, E. Bueso, J. Forment, R. Serrano and I. Kranner (2022). Comparative analysis of wild-type accessions reveals novel determinants of Arabidopsis seed longevity. Plant, cell & environment, 45(9), 2708–2728.

Nishizawa-Yokoi, A., Y. Yabuta and S. Shigeoka (2008). The contribution of carbohydrates including raffinose family oligosaccharides and sugar alcohols to protection of plant cells from oxidative damage. Plant signaling & behavior, 3(11), 1016–1018.

Oñate-Sánchez, L. and J. Vicente-Carbajosa (2008). DNA-free RNA isolation protocols for Arabidopsis thaliana, including seeds and siliques. BMC research notes, 1(1), 93.

Pascual, L. S., C. Segarra-Medina, A. Gómez-Cadenas, M. F. López-Climent, V. Vives-Peris and S. I. Zandalinas (2022). Climate change-associated multifactorial stress combination: A present challenge for our ecosystems. Journal of Plant Physiology, 276, 153764.

Pellizzaro, A., M. Neveu, D. Lalanne, B. Ly Vu, Y. Kanno, M. Seo, O. Leprince and J. Buitink (2020). A role for auxin signaling in the acquisition of longevity during seed maturation. New Phytologist, 225(1), 284–296.

Penfield, S. and D. R. MacGregor (2017). Effects of environmental variation during seed production on seed dormancy and germination. Journal of experimental botany, 68(4), 819–825.

Prieto-Dapena, P., R. Castano, C. Almoguera and J. Jordano (2006). Improved resistance to controlled deterioration in transgenic seeds. Plant physiology, 142(3), 1102–1112.

Rajjou, L. and I. Debeaujon (2008). Seed longevity: survival and maintenance of high germination ability of dry seeds. Comptes rendus biologies, 331(10), 796–805.

Renard, J., R. Niñoles, I., Martínez-Almonacid, B., Gayubas, R., Mateos-Fernández, G., Bissoli, E., Bueso, R. Serrano and J. Gadea (2020). Identification of novel seed longevity genes related to oxidative stress and seed coat by genome-wide association studies and reverse genetics. Plant, cell & environment, 43(10), 2523–2539.

Righetti, K., J. L. Vu, S. Pelletier, B. L. Vu, E. Glaab, D. Lalanne, A. Pasha, R. V. Patel, N. J. Provart and J. Verdier (2015). Inference of longevity-related genes from a robust coexpression network of seed maturation identifies regulators linking seed storability to biotic defense-related pathways. The plant cell, 27(10), 2692–2708.

Roessner, U., C. Wagner, J. Kopka, R. N. Trethewey and L. Willmitzer (2000). Simultaneous analysis of metabolites in potato tuber by gas chromatography–mass spectrometry. the plant journal, 23(1), 131–142.

Salvi, P., V. Varshney and M. Majee (2022). Raffinose family oligosaccharides (RFOs): role in seed vigor and longevity. Bioscience Reports, 42(10).

Schmuths, H., K. Bachmann, W. E. Weber, R. Horres and M. H. Hoffmann (2006). Effects of preconditioning and temperature during germination of 73 natural accessions of Arabidopsis thaliana. Annals of botany, 97(4), 623–634.

Soengas, P., V. M. Rodríguez, P. Velasco and M. E. Cartea (2018). Effect of temperature stress on antioxidant defenses in Brassica oleracea. ACS omega, 3(5), 5237–5243.

Springthorpe, V. and S. Penfield (2015). Flowering time and seed dormancy control use external coincidence to generate life history strategy. elife, 4, e05557.

Sun, M., S. Sun, C. Mao, H. Zhang, C. Ou, Z. Jia, Y. Wang, W. Ma, M. Li and S. Jia (2022). Dynamic responses of antioxidant and glyoxalase systems to seed aging based on full-length transcriptome in oat (Avena sativa L.). Antioxidants, 11(2), 395.

Sun, S., C. Mi, W. Ma and P. Mao (2024). Dynamic responses of germination characteristics and antioxidant systems to alfalfa (Medicago sativa) seed aging based on transcriptome. Plant Physiology and Biochemistry, 217, 109205.

Supek F, Bošnjak M, Škunca N, Šmuc T. REVIGO summarizes and visualizes long lists of gene ontology terms. PLoS One. 2011;6(7):e21800

Vidigal, D. S., A. C. Marques, L. A. Willems, G. Buijs, B. Méndez-Vigo, H. W. Hilhorst, L. Bentsink, F. X. Picó and C. Alonso-Blanco (2016). Altitudinal and climatic associations of seed dormancy and flowering traits evidence adaptation of annual life cycle timing in Arabidopsis thaliana. Plant, cell & environment, 39(8), 1737–1748.

Walters, R. G. (2005). Towards an understanding of photosynthetic acclimation. Journal of experimental botany, 56(411), 435–447.

Wang, B., R. Yang, Z. Zhang, S. Huang, Z. Ji, W. Zheng, H. Zhang, Y. Zhang and F. Feng (2023). Integration of miRNA and mRNA analysis reveals the role of ribosome in to anti-artificial aging in sweetcorn. International Journal of Biological Macromolecules, 240, 124434.

Wang, J., N. Nan, L. Shi, N. Li, S. Huang, A. Zhang, Y. Liu, P. Guo, B. Liu and Z. Y. Xu (2020). Arabidopsis BRCA1 represses RRTF1-mediated ROS production and ROS-responsive gene expression under dehydration stress. New Phytologist, 228(5), 1591–1610.

Wang, X., Y. Zhu, L. Tang, Y. Wang, R. Sun and X. Deng (2024). Arabidopsis HSFA9 acts as a regulator of heat response gene expression and the acquisition of thermotolerance and seed longevity. Plant and Cell Physiology, 65(3), 372–389.

Weston, E., K. Thorogood, G. Vinti and E. López-Juez (2000). Light quantity controls leaf-cell and chloroplast development in Arabidopsis thaliana wild type and blue-light-perception mutants. Planta, 211(6), 807–815.

Wozny, D., K. Kramer, I. Finkemeier, I. F. Acosta and M. Koornneef (2018). Genes for seed longevity in barley identified by genomic analysis on near isogenic lines. Plant, Cell & Environment, 41(8), 1895–1911.

Yin, G., X. Xin, C. Song, X. Chen, J. Zhang, S. Wu, R. Li, X. Liu and X. Lu (2014). Activity levels and expression of antioxidant enzymes in the ascorbate–glutathione cycle in artificially aged rice seed. Plant Physiology and Biochemistry, 80, 1–9.

Zandalinas, S. I., F. B. Fritschi and R. Mittler (2020). Signal transduction networks during stress combination. Journal of Experimental Botany, 71(5), 1734–1741.

Zandalinas, S. I. and R. Mittler (2022). Plant responses to multifactorial stress combination. New Phytologist, 234(4), 1161–1167.

Zhang, K., Y. Zhang, J. Sun, J. Meng and J. Tao (2021). Deterioration of orthodox seeds during ageing: Influencing factors, physiological alterations and the role of reactive oxygen species. Plant Physiology and Biochemistry, 158, 475–485.

Zhang, Y., D. Li, L. M. Dirk, A. B. Downie and T. Zhao (2021). ZmAGA1 hydrolyzes RFOs late during the lag phase of seed germination, shifting sugar metabolism toward seed germination over seed aging tolerance. Journal of Agricultural and Food Chemistry, 69(39), 11606–11615.

Zhao, C., B. Liu, S. Piao, X. Wang, D. B. Lobell, Y. Huang, M. Huang, Y. Yao, S. Bassu and P. Ciais (2017). Temperature increase reduces global yields of major crops in four independent estimates. Proceedings of the National Academy of sciences, 114(35), 9326–9331.

Zinsmeister, J., D. Lalanne, E. Terrasson, E. Chatelain, C. Vandecasteele, B. L. Vu, C. Dubois-Laurent, E. Geoffriau, C. L. Signor and M. Dalmais (2016). ABI5 is a regulator of seed maturation and longevity in legumes. The Plant Cell, 28(11), 2735–2754.

Zinsmeister, J., O. Leprince and J. Buitink (2020). Molecular and environmental factors regulating seed longevity. Biochemical Journal, 477(2), 305–323.

