## Supplemental Figures for "Maternal Light and Temperature Modulate Seed Longevity in Arabidopsis"

### Slide 1
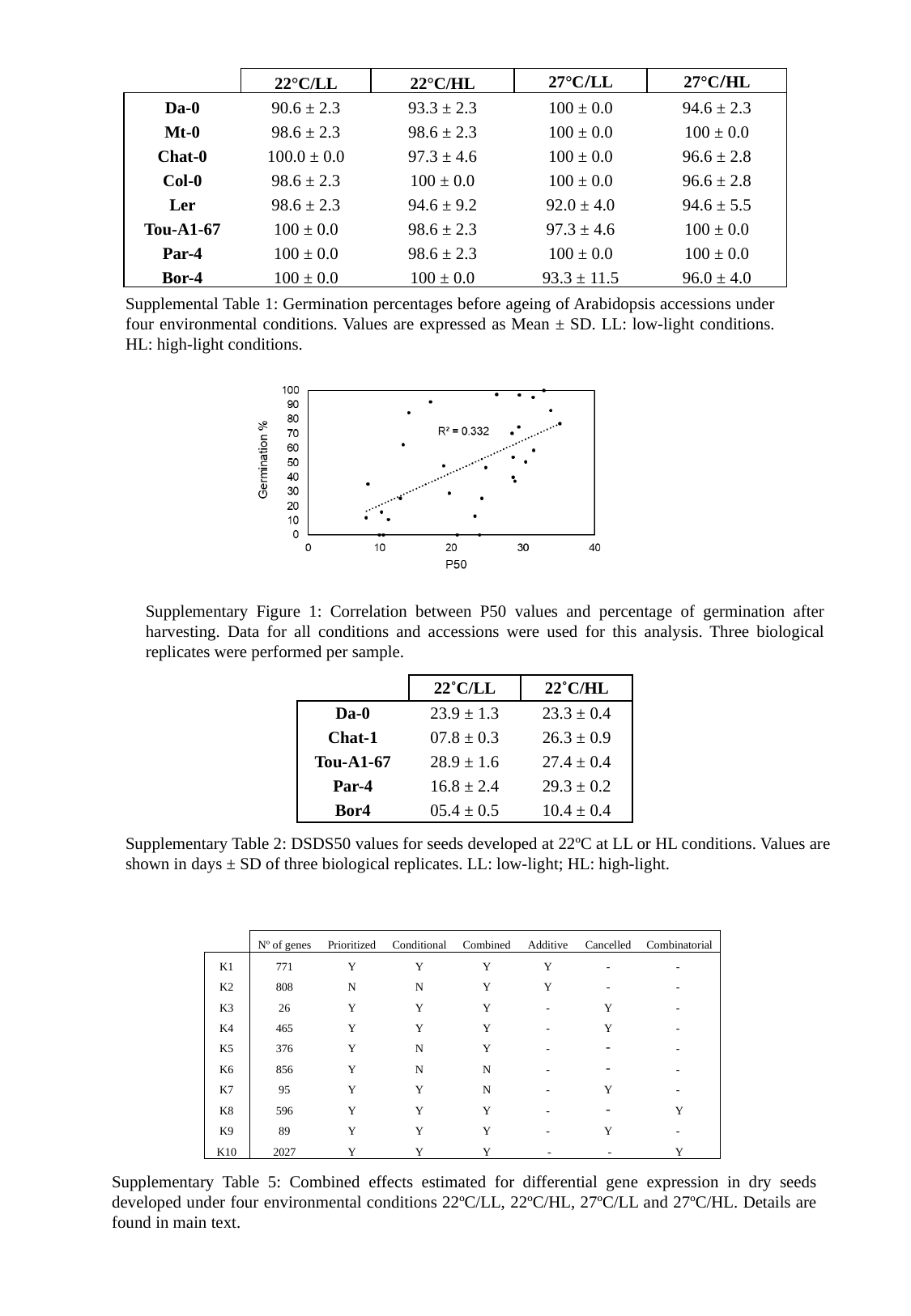

| | 22°C/LL | 22°C/HL | 27°C/LL | 27°C/HL |
| --- | --- | --- | --- | --- |
| Da-0 | 90.6 ± 2.3 | 93.3 ± 2.3 | 100 ± 0.0 | 94.6 ± 2.3 |
| Mt-0 | 98.6 ± 2.3 | 98.6 ± 2.3 | 100 ± 0.0 | 100 ± 0.0 |
| Chat-0 | 100.0 ± 0.0 | 97.3 ± 4.6 | 100 ± 0.0 | 96.6 ± 2.8 |
| Col-0 | 98.6 ± 2.3 | 100 ± 0.0 | 100 ± 0.0 | 96.6 ± 2.8 |
| Ler | 98.6 ± 2.3 | 94.6 ± 9.2 | 92.0 ± 4.0 | 94.6 ± 5.5 |
| Tou-A1-67 | 100 ± 0.0 | 98.6 ± 2.3 | 97.3 ± 4.6 | 100 ± 0.0 |
| Par-4 | 100 ± 0.0 | 98.6 ± 2.3 | 100 ± 0.0 | 100 ± 0.0 |
| Bor-4 | 100 ± 0.0 | 100 ± 0.0 | 93.3 ± 11.5 | 96.0 ± 4.0 |
Supplemental Table 1: Germination percentages before ageing of Arabidopsis accessions under four environmental conditions. Values are expressed as Mean ± SD. LL: low-light conditions. HL: high-light conditions.
Supplementary Figure 1: Correlation between P50 values and percentage of germination after harvesting. Data for all conditions and accessions were used for this analysis. Three biological replicates were performed per sample.
| | 22˚C/LL | 22˚C/HL |
| --- | --- | --- |
| Da-0 | 23.9 ± 1.3 | 23.3 ± 0.4 |
| Chat-1 | 07.8 ± 0.3 | 26.3 ± 0.9 |
| Tou-A1-67 | 28.9 ± 1.6 | 27.4 ± 0.4 |
| Par-4 | 16.8 ± 2.4 | 29.3 ± 0.2 |
| Bor4 | 05.4 ± 0.5 | 10.4 ± 0.4 |
Supplementary Table 2: DSDS50 values for seeds developed at 22ºC at LL or HL conditions. Values are shown in days ± SD of three biological replicates. LL: low-light; HL: high-light.
Supplementary Table 5: Combined effects estimated for differential gene expression in dry seeds developed under four environmental conditions 22ºC/LL, 22ºC/HL, 27ºC/LL and 27ºC/HL. Details are found in main text.

### Slide 2
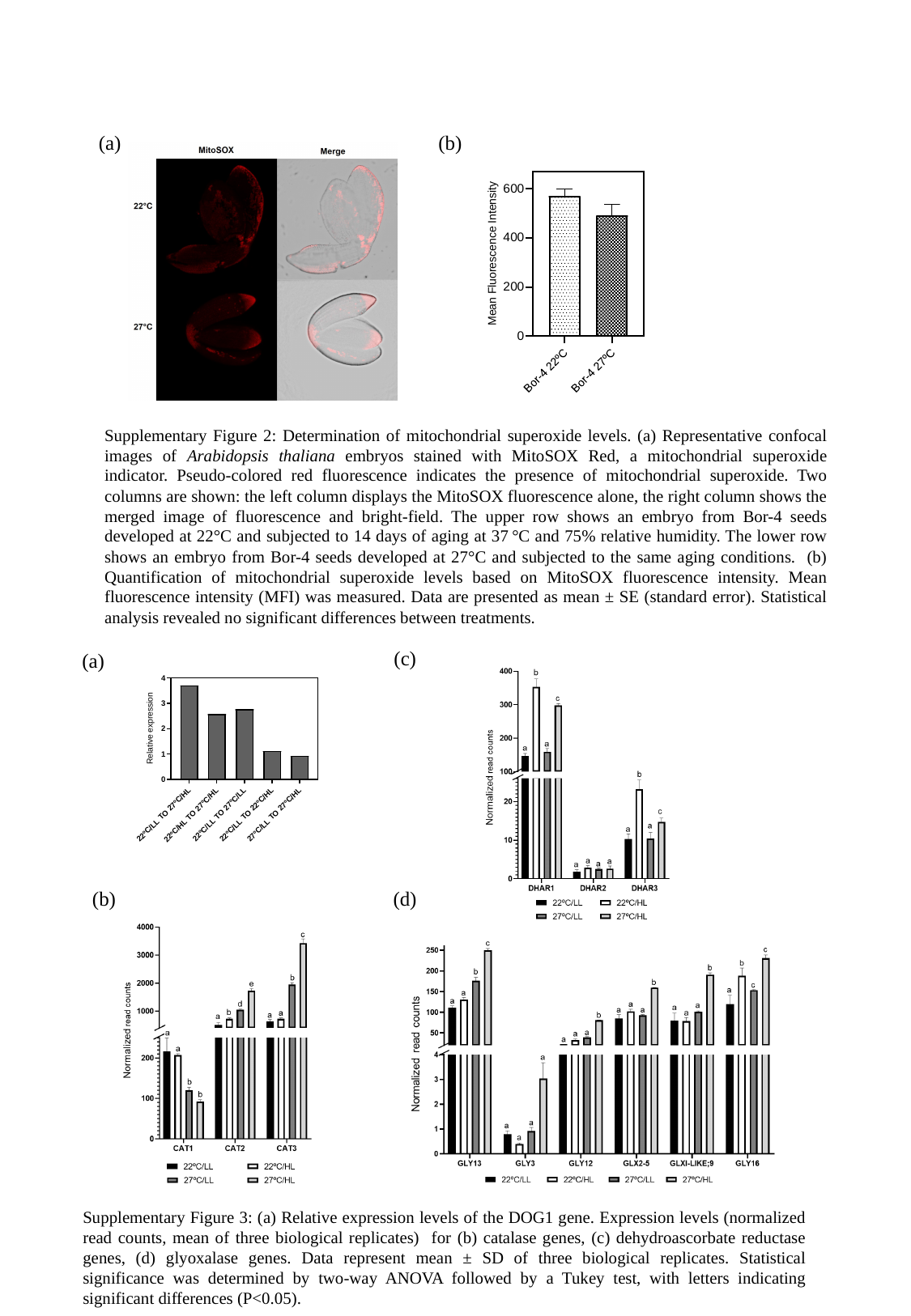

(b)
(a)
Supplementary Figure 2: Determination of mitochondrial superoxide levels. (a) Representative confocal images of Arabidopsis thaliana embryos stained with MitoSOX Red, a mitochondrial superoxide indicator. Pseudo-colored red fluorescence indicates the presence of mitochondrial superoxide. Two columns are shown: the left column displays the MitoSOX fluorescence alone, the right column shows the merged image of fluorescence and bright-field. The upper row shows an embryo from Bor-4 seeds developed at 22°C and subjected to 14 days of aging at 37 °C and 75% relative humidity. The lower row shows an embryo from Bor-4 seeds developed at 27°C and subjected to the same aging conditions. (b) Quantification of mitochondrial superoxide levels based on MitoSOX fluorescence intensity. Mean fluorescence intensity (MFI) was measured. Data are presented as mean ± SE (standard error). Statistical analysis revealed no significant differences between treatments.
(c)
(a)
(b)
(d)
Supplementary Figure 3: (a) Relative expression levels of the DOG1 gene. Expression levels (normalized read counts, mean of three biological replicates) for (b) catalase genes, (c) dehydroascorbate reductase genes, (d) glyoxalase genes. Data represent mean ± SD of three biological replicates. Statistical significance was determined by two-way ANOVA followed by a Tukey test, with letters indicating significant differences (P<0.05).
