## Supplemental Table 9 for "Maternal Light and Temperature Modulate Seed Longevity in Arabidopsis"

Supplemental Table 9. A list of primers used in this study

| Primer | Sequence (5′→ 3′) |
|---|---|
| M1080 RT FP | AAAATCAATAGGTGCCGGAGCTGC |
| M1080 RT RP | CGGTTAGAGCAAAGCCCAAAATGG |
| AT2G07698 RT-FP | CCTCGAAGAAAGAAGAGCTGCGG |
| AT2G07698 RT-RP | ACCACCAAAGACAACAATCCCGAC |
| M00640 RT-FP | GGATCAGCTTGCGAATTTGTGGC |
| M00640 RT-RP | GCAAATTGCTTCCCCACTAAGGTG |
| AT3G52300 RT-FP | GCTCCAGACCAAATTTAGTCAGGAACC |
| AT3G52300 RT-RP | CTGTTCTGCTTCTTTCAGTTCCACCAAC |
| MG00510 RT-FP | GGTGTAATGTTAAGAGGTCCAGGGG |
| MG00510 RT-RP | CACAATGATCCGAAGACTTTGTCGC |
| AT2G07687 RT-FP | TTCTTCTTTGGCACCTGCGGTAG |
| AT2G07687 RT-RP | CCCCGCGAGTATAGCATGATGAG |
| MG01360 RT-FP | TTGGTTCTTCGGTCATCCAGAGG |
| MG01360 RT-RP | TGATCATGGTAGCTGCGGTGAAG |
| AT2G07727 RT-FP | GGTGTAGCCGCAATAGCACCAG |
| AT2G07727 RT-RP | GAATGGGCGTTATGGCAAAGAACAAG |
| AT1G11840 RT-FP | TCAAATTCTATGAAAAGGCCCTCGGG |
| AT1G11840 RT-RP | CAATCTGTGCATATGCATTGCCCTTTG |
| AT1G08110 RT-FP | GTGTGCTGGGGATGTCATTGC |
| AT1G08110 RT-RP | GTCAATTCAATTGTTGCCGGTTGACC |
| AT1G08110 RT-FP | TGTACTTCCTGGGCTACGAGGATAC |
| AT1G08110 RT-RP | CGGTAACCCCAATGTGCCC |
| AT1G20620 RT FP | CTCACCATCGGAGAAAGAGGTCC |
| AT1G20620 RT RP | ACACCAGGGGCTCTGAGAAAATC |
| AT1G19570 RT-FP | GATCATCTCGGCGACTGTCCG |
| AT1G19570 RT-RP | GACTAATGTCCAAGAACCACTGGGG |
